# scDIAGRAM: Detecting Chromatin Compartments from Individual Single-Cell Hi-C Matrix without Imputation or Reference Features

**DOI:** 10.1101/2025.08.01.668129

**Authors:** Yongli Peng, Yujing Deng, Menghan Liu, Zhiyuan Liu, Ya-Hui Li, Xiang-Yu Zhao, Dong Xing, Jinzhu Jia, Hao Ge

**Affiliations:** Beijing International Center for Mathematical Research(BICMR), Peking University, 100871, China; Biomedical Pioneering Innovation Center(BIOPIC), Peking University, 100871, China; Peking University Institute of Haematology, National Clinical Research Center for Haematologic Disease, Peking University People’s Hospital, 100044, China; Beijing Advanced Innovation Center for Genomics, Peking University, 100871, China; School of Public Health and Center for Statistical Science, Peking University, 100871, China

**Keywords:** single-cell Hi-C, A/B compartment, statistical modeling, chromatin heterogeneity

## Abstract

Single-cell Hi-C (scHi-C) provides unprecedented insight into 3D genome organization, but its sparse and noisy data pose challenges in accurately detecting A/B compartments, which are crucial for understanding chromatin structure and gene regulation. We presented scDIAGRAM, a data-driven method for annotating A/B compartments in single cells using direct statistical modeling and graph community detection. Unlike existing approaches, scDIAGRAM operates without relying on external information, such as the CpG density or imputation techniques, and preserves cell-to-cell heterogeneity. Accuracy and robustness of scDIAGRAM were illustrated through simulated scHi-C datasets and a human cell line. We applied scDIAGRAM to real scHi-C datasets from the mouse brain cortex, mouse embryonic development, and human acute myeloid leukemia (AML), demonstrating its ability to capture compartmental shifts associated with transcriptional variation. This robust framework offers new insights into the functional roles of chromatin compartments at single-cell resolution across various biological contexts.

## 1 Introduction

Advancements in three-dimensional (3D) whole-genome mapping techniques, such as Hi-C, have significantly improved our understanding of genome organization within the nucleus [1, 2, 3, 4, 5]. Bulk Hi-C studies have revealed that the genome is organized into hierarchical structures, including chromosome territory [6], A/B compartments [1], topologically associating domains (TADs) [7, 8], and chromatin loops [9]. More recently, the integration of single-cell mapping technologies with conventional Hi-C has led to the emergence of single-cell Hi-C (scHi-C), allowing for the analysis of 3D genome organization at the resolution of individual cells [10, 11, 12, 13, 14].

However, scHi-C data are typically sparse and noisy, posing substantial challenges in understanding the variability of these structures at the single-cell level [15, 16, 17, 18, 19]. To address these challenges, computational tools have been developed, either by improving data quality through imputation [20, 21, 22, 23, 24, 25] or by explicitly detecting multi-scale genome structures in individual cells without imputation [26, 27, 28, 29].

In this work, we focused on annotating A/B compartments using only individual single-cell contact matrices. In bulk Hi-C analysis, compartmentalization is typically inferred from the normalized observed/expected (O/E) matrix or correlation matrices [1]. Principal component analysis (PCA) applied to the correlation matrix classifies the genome into open (A) and closed (B) compartments, corresponding to euchromatin and heterochromatin, respectively. This classification is further supported by fluorescence in situ hybridization (FISH) experiments [30, 31]. Recently, tools such as cooltools [32] have streamlined this process for bulk Hi-C data.

Several computational methods were developed to identify single-cell compartments (scCompartments) in scHi-C data, often relying on external genomic features or imputation to address data sparsity. The A compartment was typically associated with higher gene density, greater CpG density, and stronger correlations with active histone modifications [1, 12, 33, 34]. Accordingly, one approach leveraged the CpG density from a reference genome combined with scHi-C matrices to infer A/B compartments in individual cells [12]. This method, refered as scA/B values, treated the Hi-C matrix as a graph and averaged CpG values across neighboring loci. This reference-guided strategy might produce annotations that reflect CpG patterns rather than the true 3D chromatin structure of single cells. Other methods focused on imputing missing contacts in sparse scHi-C matrices before applying compartment annotation techniques originally developed for bulk Hi-C, such as PCA [21, 20, 13, 22]. Some approaches, like Higashi [21, 22], incorporated information from similar or neighboring cells to enhance imputation, which could possibly blur biological differences and lead to an averaging effect. Even within-cell imputation approaches, such as that employed by scHiCluster [20], carried the risk of distorting compartment structures due to potential errors in imputed values. These challenges highlight the need for a method that directly analyzes single-cell Hi-C data without external dependencies, preserving both the integrity of chromatin structures and the biological variability across cells.

Here, we introduced scDIAGRAM (single-cell compartments annotation by DIrect stAtistical modeling and GRAph coMmunity detection), a novel computational tool designed to annotate chromatin A/B compartments in scHi-C data. The method addressed the challenges posed by sparse and noisy scHi-C datasets by applying direct statistical modeling and graph community detection [35, 36] to group genomic loci into A/B compartments, without relying on external data or imputation. scDIAGRAM was tested on both simulated and real datasets, including mouse brain cortex, mouse developing embryos and human acute myeloid leukemia (AML) [37, 38], showing its ability to preserve compartmental heterogeneity and capture dynamic changes in genome organization. By linking compartmental shifts to transcriptional and epigenetic variations, sc-DIAGRAM enhanced our understanding of the functional roles of chromatin compartments and offered a robust framework for exploring 3D genome organization at single-cell resolution across diverse biological systems.

## 2 Methods

scDIAGRAM took an intrachromosomal Hi-C contact matrix as input, in which the genome was divided into discrete regions (i.e., “loci” or “bin”) at a specified resolution. The input data could be either bulk Hi-C or scHi-C matrices, with varying resolutions. The workflow began with 2D change-point detection, followed by the step of graph partitioning, to address the significant noise inherent in scHi-C data and annotate compartments for each loci. The underlying computational workflow was summarized in Fig. 1. Further details of the algorithm were provided below.

**Figure 1:**
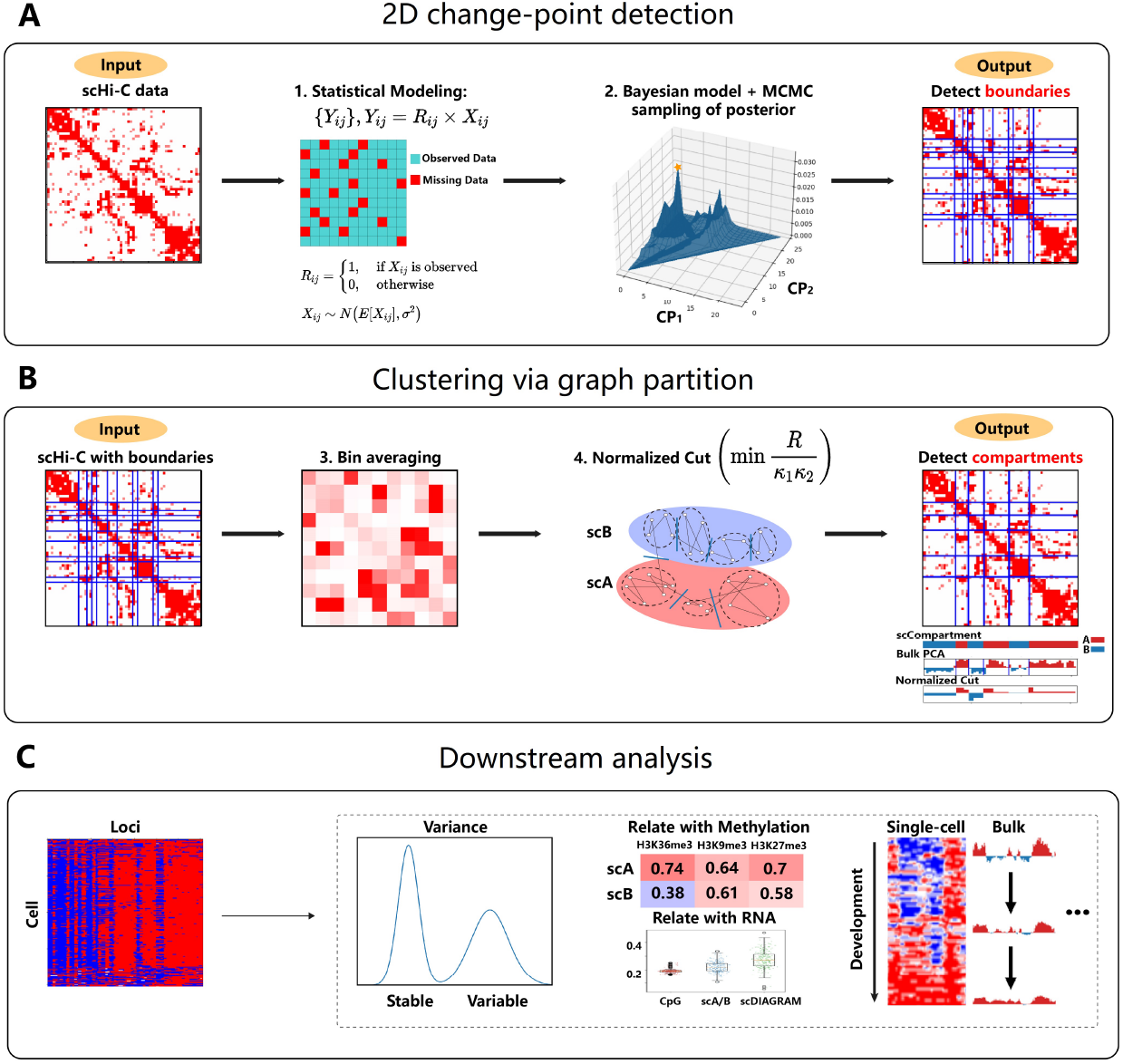
Schematic workflow of scDIAGRAM. (A). scDIAGRAM takes a scHi-C matrix as input, treating the problem as a 2D change-point (CP) detection problem. A direct statistical model is established, and scDIAGRAM employs a Bayesian model with Markov Chain Monte Carlo (MCMC) to detect CP positions. (B). Bin averaging is performed to each block formed by the detected CPs, and then a graph partitioning algorithm is applied to classify the genomic loci groups separated by CPs into two compartments. (C) The output is a heatmap representing the annotated compartments for each genomic loci. Each row corresponds to a cell, and each column represents a loci. The heatmap can be binary (A or B compartments) or real-valued (Ncut values), with red denoting the A compartment, and higher Ncut values indicating more active loci. Using scDIAGRAM, one can study compartmental heterogeneity within single cells, stable and variable genomic loci, the relation between single-cell compartments with methylation and RNA expression, or analyze dynamic compartmental changes during development, among many other applications.

### 2D change-point detection

We first constructed a direct statistical model for the contact matrix and framed the annotation as a 2D change-point detection problem. We then applied Bayesian modeling and Markov Chain Monte Carlo (MCMC) methods to obtain the maximum likelihood estimation (MLE) of the problem.

For each measured contact matrix *{Y*_*ij*_*}*, in which *Y*_*ij*_ is the measured contact number between the *i*-th and *j*-th loci, we assumed that *Y*_*ij*_ = *R*_*ij*_ *× X*_*ij*_, where *R*_*ij*_ represented the dropout effect and *X*_*ij*_ denoted the true contact number inside the cell. These variables were assumed to be independent of each other.

The distributions of *X*_*ij*_ is assumed to be Gaussian, i.e. *X*_*ij*_ ∼ *N* (*E*[*X*_*ij*_], *σ*^2^), and *R*_*ij*_ follows a Bernoulli distribution. If *X*_*ij*_ is observed, *R*_*ij*_ = 1; and it’s 0 otherwise.

We focused on detecting the compartment boundaries as change-points (CPs) in the 2D contact matrix. Specifically, *K* CPs divide the matrix into (*K* + 1) × (*K* + 1) blocks. We assumed that the parameters *P* (*R*_*ij*_ = 1) and *E*[*X*_*ij*_] were constant for all pairs (*i, j*) within the same block. These parameters were represented by two (*K* + 1) *×* (*K* + 1) matrices, denoted as 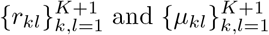. We could solve the 2D change-point detection problem using maximum likelihood estimation (MLE).

Since each gap between adjacent loci could be considered as a CP, explicitly computing the MLE was computationally infeasible. Therefore, we turned to a Bayesian model and employed MCMC sampling to obtain samples of the parameters and then obtained the MLE.

First, we used one-hot encoding for the CP positions. We assumed that the dimension of contact matrix was *n*, which resulted in (*n* −1) potential CP positions in total. The state space for CP positions was {0, 1}^*n*−1^, where 1 indicated the presence of a CP at a given position and 0 otherwise.

Next, we imposed priors on these states. We set *K* CPs in total (*K* is a hyperparameter and shall be determined in advance), then the prior was 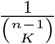 (uniform across states) and the posterior was proportional to the likelihood. In this way, the genomic loci 1, 2, *‥, n* were divided into *K* + 1 groups, denoted as *g*_1_, *g*_2_, *‥, g*_*K*+1_.

By Bayes’ formula, we had the posterior distribution

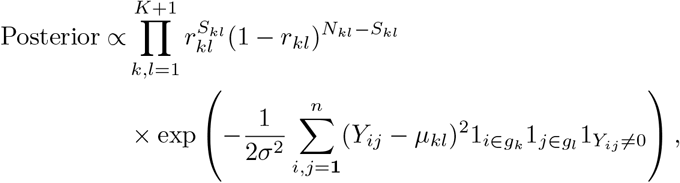

where *N*_*kl*_ was the sample size of the (*k, l*)-block and 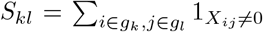 was the nonzero sample size in the (*k, l*)-block. *σ*^2^, *µ*_*kl*_, and *r*_*kl*_ needed to be estimated from the data once the positions of all CPs were given. For simplicity, we used the sample variance and the sample mean.

Thus

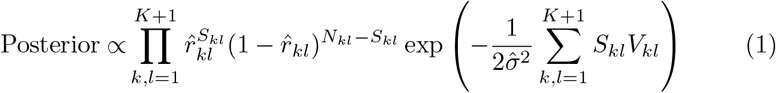

Where 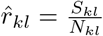, and *V*_*kl*_ was the sample variance of nonzero entries inside the (*k, l*)-block.

We used Metropolis-Hasting (MH) methods to sample from the posterior distribution (or just the likelihood). See Supplementary Methods for details of the algorithm.

### Clustering of loci group via graph partition

With the positions of all *K* CPs identified, all genomic loci were divided into (*K* + 1) groups. We employed a graph partitioning approach to cluster these groups into two compartments.

We treated each group of loci (bins) between two adjacent CPs as a node in a graph. The *K* CPs divided the matrix into (*K* + 1) × (*K* + 1) blocks. We computed the average value of contact numbers within each block, as the weight for the edge connecting the two nodes, thereby reducing the original large *n* ×*n* matrix into a smaller (*K* + 1) × (*K* + 1) one.

We applied a graph partitioning framework called normalized cut to the constructed weighted graph [35, 36], for its simplicity and effectiveness. It was formulated as an optimization problem min 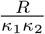, where *R* represented the graph cut (the total number of contacts between A and B compartments) and *κ*^1^, *κ*^2^ were the sums of degrees in the two clustered compartments.

This problem can be addressed using spectral methods. The second largest eigenvector *ν*_2_ of **D**^−1*/*2^**AD**^−1*/*2^ is exactly a relaxed solution, in which **A** is the adjacency matrix and **D** is the diagonal matrix with elements *D*_*ii*_ = *k*_*i*_. *k*_*i*_ is the degree of node *i*. See Supplementary Methods for more details.

To obtain a discrete-valued vector for graph partitioning, *ν*_2_ was rounded, typically using 0 as a threshold and nodes were assigned to the two compartments based on the sign of *ν*_2_ (according to [35], the result was robust to the rounding strategy chosen for division). The compartmental value of each locus just inherited from the values of the corresponding node, i.e. the group of loci separated by CPs.

The remaining step is to add A/B annotation, i.e. determining which compartment is A or B. Based on bulk Hi-C analysis, we knew that the CpG density in the reference genome was strongly correlated with compartments, with the A compartment typically corresponding to regions of high CpG density. Following the convention in bulk Hi-C studies, we determined the sign by comparing the Pearson correlation between *ν*_2_ with the CpG density. A low absolute value of the correlation may indicate weak compartmentalization shown at this single cell.

### Hyperparameter settings

The number of CPs was determined by comparing A/B compartment annotations generated by scDIAGRAM at increasing CP numbers. Specifically, we assessed the intersection (defined later) of these annotations and selected the CP number at which increasing this value no longer resulted in significant changes to the A/B annotations. For scHi-C datasets, we first determined the CP number for several cells and obtained the largest one among them. Then this CP number was fixed and applied to all other cells in this dataset. Typically, this value was set much higher than the CP number used for bulk Hi-C data. The choice of CP number also depended on the resolution; for example, a reasonable choice was approximately 100 CPs at the resolution of 100 kilobase (kb) and 2040 CPs at 1 megabase (Mb). In this study, we mainly focused on scHi-C matrices at 100 kb resolution. Comparable results could also be obtained at 1 Mb resolution, albeit with coarser structural details. We tested scDIAGRAM on a real single cell and found that the results remained stable as the fixed number of CPs increased (Supplementary Figure S1).

### Simulated scHi-C matrices

To benchmark the accuracy and robustness of scDIAGRAM, we generated simulated single-cell Hi-C datasets derived from imaging-based 3D structural models, bulk Hi-C matrices, and high-coverage single-cell data. These datasets were designed to reflect varying levels of sparsity and biological heterogeneity observed in real data. Details of the simulation protocols, parameter settings, and ground-truth annotations are provided in the Supplementary Methods.

### Real scHi-C and scRNA-seq datasets

Publicly available scHi-C datasets from mouse brain and mouse embryonic development were processed using standard pipelines. Contact matrices were normalized using cooler and cooltools, and pseudo-bulk matrices were constructed for each condition or cell type. We ran scDIAGRAM on each chromosome separately.

We collected acute myeloid leukemia (AML) patient samples and their pathological results from the Department of Hematology at Peking University People’s Hospital. All samples were obtained via bone marrow aspiration. The study was approved by the Ethics Committee of Peking University People’s Hospital (2024PHB391-001). All patients signed informed consent forms as required. The data processing method for AML in this study follows the same procedure as in [37].

ScRNA-seq data were preprocessed with Seurat for clustering and marker genes were identified using Seurat [39] with default parameters.

See Supplementary Methods for more details on the real single-cell datasets.

### Comparing scDIAGRAM with other methods

We compared scDIAGRAM with scA/B values [12], compartments derived from imputed scHi-C matrices, and from 3D structure modeling.

The output of scA/B is only a real valued compartmentalization. To obtain binary annotations, we applied rank/quantile normalization to transform these values into the range [0,1], and then binarized the compartments using a cutoff of 1/2. We used scHiCluster with default parameters [20] to impute the scHi-C matrices, then applied cooltools [32] on the imputed matrices to generate the scCompartments.

We trained Higashi without using neighboring cell information (0 nbr) [21] to impute the scHi-C matrices and applied its built-in method to call compartments at the single-cell level. Other parameters were set as default.

We also utilized the 3D modeling algorithm by the Hickit software with default parameters [12], on downsampled scHi-C datasets. Once the structure was generated, we took the inverse of the pairwise spatial distance matrices and applied cooltools to them. In this manner, we treated the 3D modeling as another method of imputing scHi-C matrices.

#### 2.0.1 Evaluation criteria

The methods were evaluated using several metrics, including correlation, intersection, and accuracy. Both Pearson and Spearman correlations were accessed. Intersection was quantified as the fraction of correctly assigned loci into A/B compartments for each chromosome in each cell. Accuracy at each loci was similarly defined as the fraction of correctly assigned samples among all samples, again focusing on loci classification.

### 2.1 Stable and variable single-cell compartments

We computed the variance of binary compartments (1 for A and 0 for B) for each loci across all cells, with values ranging from 0 to 0.25. Genomic loci with stable and variable compartments were identified by applying a cutoff at the 50th percentile of this variance [24, 40]. To further explore the robustness of this classification, we also examined the top and bottom 25th, 10th, and 5th percentiles of the variance.

## 3 Results

### 3.1 scDIAGRAM is validated on simulated scHi-C matrices and a human cell line

We began by validating the performance of scDIAGRAM in comparison to scA/B and compartments obtained from scHiCluster and Higashi imputation, using simulated scHi-C datasets and scHi-C data from a human cell line.

#### 3.1.1 Simulation via downsampling pseudo-bulk Hi-C data

We first generated synthetic data through downsampling a pseudo-bulk Hi-C matrix of the Ex1 cell type on chr7 [37], with sample rates ranging from 1/400 to 1/3200 (Supplementary Methods). First of all, scDIAGRAM accurately predicted the correct compartments in the pseudo-bulk data itself, in comparison to that obtained using cooltools, with 0.94 intersection and 0.84 Pearson correlation. scA/B only had 0.825 intersection and 0.6 Pearson correlation (Supplementary Figures S2).

For downsampled data, scDIAGRAM showed closer alignment with the ground truth than scA/B, both for binary compartments (measured by intersection) and real-valued compartments (measured by Spearman correlation), across various sample rates (Fig. 2A, Supplementary Figure S3A, B). Similar results were observed for chromosome 7 of the mixed late mesenchyme cell type (Supplementary Figure S3C).

**Figure 2:**
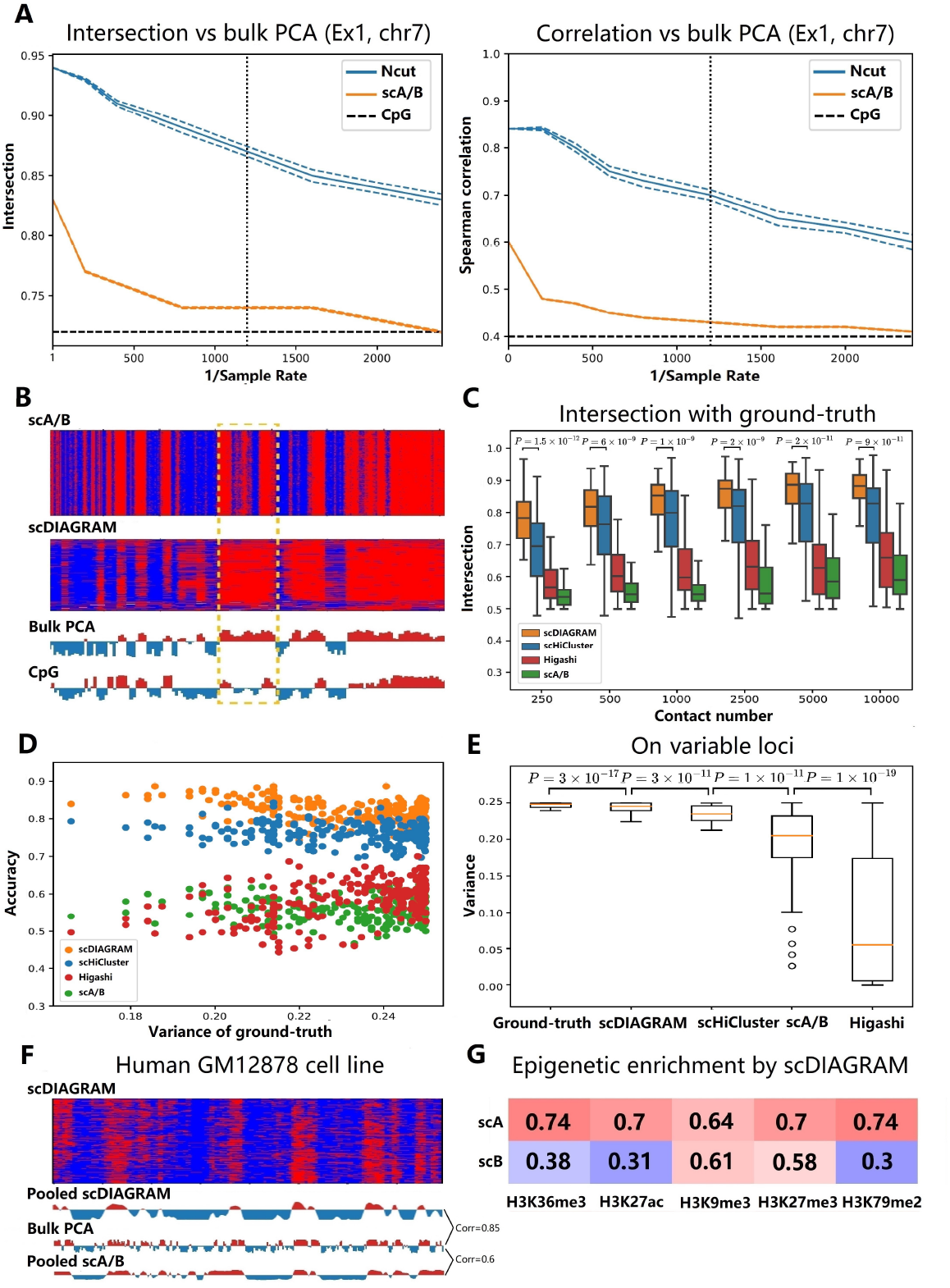
Evaluation of the performance of scDIAGRAM in simulated scHi-C dataset and a human cell line GM12878. (A). The intersection and Spearman correlation versus bulk PCA on a simulated scHi-C dataset via downsampling a pseudo-bulk Hi-C matrix. The vertical dash line is the sample rate of a typical real scHi-C dataset, and the horizontal dash line is the intersection or correlation between the CpG density and bulk PCA. (B). scDIAGRAM is more close to bulk PCA compared to scA/B, illustrated by the heatmap from the same downsampled data as (A). Each row represents a cell. As indicated in the boxed region, the CpG density is quite different from bulk PCA. (C and D). The intersection and accuracy on a simulated scHi-C dataset via downsampling a single-cell 3D imaging data. Accuracy is computed when the contact number is 1000, with genomic loci ordered by its compartmental variance calculated from ground-truth. The p-value *P* = 7 × 10^−82^ between the accuracy of scDIAGRAM and scHiCluster in (D). (E). scDIAGRAM is more heterogeneous compared with scHiCluster, Higashi and scA/B. (F). Heatmap of scDIAGRAM and bulk PCA on human cell line GM12878. (G). Enrichment of epigenomic signals (average of these signals’ fold change) in pseudo-bulk compartments.

Moreover, at very low sample rates, where the data became highly sparse, scA/B often converged to the CpG signal, which is a universal feature independent of scHi-C data. In contrast, scDIAGRAM maintained better alignment with the pseudo-bulk data used to generate the simulated scHi-C matrices (Fig. 2A). This phenomenon was further highlighted by examining regions where the CpG signal and bulk PCA differed significantly (Fig. 2B).

#### 3.1.2 Simulation via downsampling single-cell 3D genome imaging data

Next, we generated synthetic data by downsampling single-cell 3D genome imaging data [41], with contact numbers ranging from 250 to 10,000, which is comparable to real scHi-C datasets (Supplementary Methods, Supplementary Figure S4). We used chromosome 2 and a dataset of 300 cells, similar to what have been done in [21].

We compared the performance of scDIAGRAM with scA/B, scHiCluster followed by PCA and Higashi with its built-in compartments (Fig. 2C, Supplementary Figure S5A). scDIAGRAM outperformed scHiCluster in binary compartment annotation (measured by intersection) (Fig. 2C), while showing comparable performance in predicting real-valued compartments (measured by Pearson and Spearman correlation; Supplementary Figure S5B). Additionally, we observed that the intersection and correlation from scHiCluster often exhibited larger variances, suggesting potential instability in the imputation. In contrast, scA/B performed poorly on downsampled data, both for binary and real-valued compartments (Fig. 2C).

Higashi performed slightly better than scA/B, but remained inferior to sc-DIAGRAM and scHiCluster. This discrepancy may stem from the pronounced heterogeneity inherent in the dataset. Higashi called single-cell compartments by projecting imputed matrices onto the pseudo-bulk compartment. As a result, it tends to produce relatively homogeneous compartments across cells (see below), suggesting that Higashi may be less suitable for datasets characterized by high heterogeneity.

With a fixed contact number, we evaluated the classification accuracy for each locus across different cells (Methods). In Fig. 2D, when the contact number was 1000, scDIAGRAM outperformed scHiCluster, Higashi and scA/B in terms of accuracy, with scA/B showing the poorest performance. Each dot in the plot represents a genomic locus, ordered by the compartmental variance of the ground truth along the x-axis. This observation held universally for loci with different variances. The same results were observed for different contact numbers (Supplementary Figure S5C).

Since 3D imaging data can generate a complete Hi-C matrix for each single cell, we were able to explore compartmental heterogeneity within this dataset. We computed the variance of binary scCompartment values across cells for each genomic loci and found that scDIAGRAM exhibited greater heterogeneity compared to scHiCluster, Higashi and scA/B (Supplementary Figure S5D), and was closer to the true heterogeneity directly obtained from the 3D imaging data. This difference was especially pronounced for the more variable loci (Fig. 2E).

#### 3.1.3 scDIAGRAM applied on the human GM12878 cell line

We applied scDIAGRAM to scHi-C data from the human GM12878 cell line at a 1 Mb resolution [13] (Fig. 2F). Compared to scA/B, the pooled scCompartments from scDIAGRAM showed a stronger Pearson correlation with bulk PCA (Correlation = 0.85, compared to scA/B’s correlation of 0.6). Additionally, genomic regions within the same compartment interacted more frequently than those in different compartments, with A − A>A − B and B − B>A−B interactions (Supplementary Figure S6).

Moreover we found these scCompartments effectively stratified histone modifications, consistent with observations from bulk PCA. We compared an epigenomic mark enrichment profile with pooled scCompartments (Fig. 2G). The pattern of histone mark enrichment in A/B compartments mirrored the observations from bulk Hi-C [1, 9]. While all histone marks tested were enriched in A compartments, activating marks like H3K36me3 showed a larger enrichment difference between A and B compartments (and a higher Pearson correlation with pseudo-bulk compartments). In contrast, repressive marks such as H3K27me3 and H3K9me3 displayed smaller enrichment differences (and lower correlation with pseudo-bulk compartments), indicating that H3K27me3 and H3K9me3 were more likely to localize to B compartments compared to other marks.

### 3.2 scDIAGRAM applied on the scHi-C data of adult mouse brain cortex

We applied scDIAGRAM to the HiRES dataset from mouse brains [37]. Embedding single cells based on scDIAGRAM-generated compartments revealed major brain cell types, which were similar to those identified using scA/B, scHiCluster or Higashi (Supplementary Figure S7A). This suggested that scCompartments alone can effectively distinguish different major cell types.

Similar behaviors observed with scDIAGRAM in simulated data were also evident in this real scHi-C dataset. For the Ex1 cell type (chr7, 100 kb), as shown in Fig. 3A, focusing on regions where the CpG signal distinctly differed from bulk PCA, we found that scDIAGRAM was more closely aligned with bulk PCA, whereas scA/B was more reflective of the CpG signal. In Fig. 3B, which considers the entire dataset (chr7, 100 kb), scDIAGRAM displayed greater heterogeneity. We computed the variance of scCompartments across cells for sc-DIAGRAM, scHiCluster, Higashi and scA/B, and for bulk PCA, we calculated the variance from bulk compartments across seven cell types. Each violin plot of variances from bulk PCA, scDIAGRAM, scA/B and Higashi exhibited two peaks, indicating stable and variable genomic regions (Fig. 3B). In contrast, scHiCluster showed large variance for all genomic loci, suggesting potential instability during imputation. The bimodal pattern was less pronounced in scA/B and Higashi, demonstrating that scA/B and Higashi typically generated more homogeneous compartments. Furthermore, when we selected the top 10% most variable loci from real-valued bulk PCA, scDIAGRAM exhibited even greater heterogeneity compared to scHiCluster, scA/B and Higashi (Fig. 3B). We also compared the pooled scCompartments generated by scDIAGRAM across Ex1 cells with the bulk PCA. The pooled scCompartments always generated better Pearson correlation than a single scCompartment, and scDIAGRAM had better Pearson correlation with bulk PCA than scA/B (Supplementary Figure S7B).

**Figure 3:**
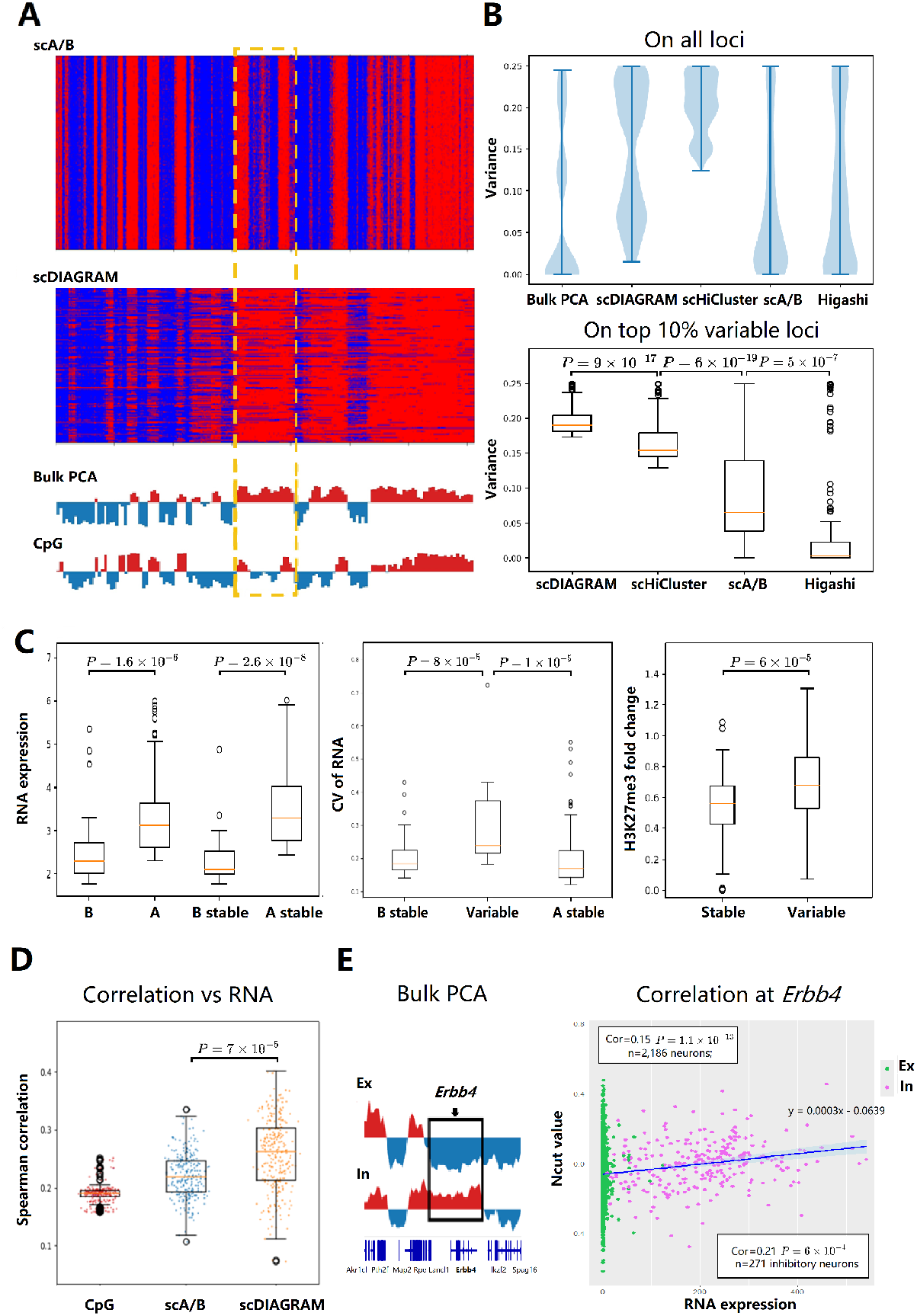
scDIAGRAM applied in a real scHi-C dataset from adult mouse brain cortex. (A). The scCompartments generated by scDIAGRAM is more close to bulk PCA than those generated by scA/B, on the Ex1 dataset from HiRES. Each row represents a cell. The highlighted region is where the CpG density is quite different from bulk PCA (B). scDIAGRAM is more heterogeneous compared with scHiCluster, Higashi and scA/B. (C). Joint analysis of single-cell transcription and compartments in the same cell type. (D). Spearman correlations with RNA expressions for CpG density, scA/B and scDIAGRAM, in single cells. (E). Visualization of the bulk PCA at the Erbb4 locus between excitatory and inhibitory neurons. The RNA expression and real-valued scCompartments (Ncut values from scDIAGRAM) for each single cell, at the Erbb4 locus. Inhibitory neurons exhibited higher Pearson correlation between RNA and scCompartments than putting inhibitory and excitatory neurons together.

One limitation of scA/B is that it generates smaller compartments, which may not reflect biologically meaningful structures, as compartments typically span multiple megabases. These smaller compartments might correspond to TAD domains or subcompartments, potentially complicating compartment annotations. In contrast, scDIAGRAM produced compartments that were more consistent in size with those from bulk PCA, while both scA/B and scHiCluster tended to generate smaller compartments (Supplementary Figure S8).

#### 3.2.1 Stable and variable single-cell compartments from scDIAGRAM

We applied scDIAGRAM to identify genomic loci with stable and variable single-cell compartments (Methods). Overall, a larger proportion of loci were stable, while variable loci predominantly corresponded to compartmental boundaries and cell-type-specific patterns of genome organization (Supplementary Figure S9A-C). For each genomic locus, classified as either variable or stable, we calculated the fraction of cells in the A and B compartments. Stable regions were predominantly annotated as A on chromosome 7 and as B on chromosome 1, whereas variable regions exhibited a more balanced distribution between the A and B compartments (Supplementary Figure S9D, E). These results are consistent with prior researches [37, 40, 42].

In the HiRES dataset, which included both Hi-C and RNA-seq data measured simultaneously for each cell, we were able to correlate the scCompartments identified by scDIAGRAM with RNA expressions, exploring their functional implications. We again focused on the Ex1 cell type at 100 kb resolution. As shown in Fig. 3C, scCompartments associated with higher RNA expression were more active. Stable loci were further categorized into A stable and B stable regions. We found that A stable loci exhibited significantly higher RNA expression compared to B stable loci.

In a manner similar to the analysis of imaging data [41], we examined changes in scCompartments for genes in transcribing (UMIs*>*10) versus silencing (UMIs*<*1) states within a single cell type (Ex1, Supplementary Figure S10A). We found that for nearly 60% of the genes studied, the compartment at their TSS was more active during transcription than when the gene was silenced. Both scDIAGRAM and scA/B produced similar patterns, with scDIAGRAM showing a larger fold change in compartmentalization, suggesting greater heterogeneity (Supplementary Figure S10A). Regions with variable compartments exhibited greater transcriptional variability (Fig. 3C), with this difference becoming more pronounced when more stringent thresholds were applied (Supplementary Figure S10B). Here, transcriptional variability was quantified by the coefficient of variation.

Similarly, we also divided genomic loci of GM12878 cells into variable and stable single-cell compartments and assessed the enrichment of H3K27me3, a histone mark that is enriched in subcompartment B1 (which is unstable) and indicative of facultative heterochromatin at the bulk level [43]. Facultative heterochromatin is a structure that can adopt either open or compact conformations depending on temporal and spatial contexts [9, 43], making it inherently unstable.

We observed significantly higher enrichment of H3K27me3 in variable genomic regions compared to stable regions in the cell population (*P* = 6 × 10^−5^, Fig. 3C). This result is further supported by the known association between H3K27me3-repressed genes and expression heterogeneity [44]. Moreover, we found a moderate Pearson correlation between the variance of scCompartments and the H3K27me3 signal (Pearson Correlation = 0.4, Supplementary Figure S10C).

#### 3.2.2 Comparing scCompartments between two cell types

We compared compartmentalization and transcription between Ex1 and astrocyte (Ast) cell types in the HiRES dataset. Comparing top 100 up and down regulated genes in Ex1, we observed elevated pseudo-bulk compartment values for up-regulated genes compared to down-regulated genes (Supplementary Figure S11A). Conversely, the top 100 compartmental activated genes in Ex1 also exhibited higher transcriptional activity than inactivated genes (Supplementary Figure S11B). The RNA fold change with all marker genes was correlated with compartmental differences, with Pearson and Spearman correlations typically ranging from 0.15 to 0.3 on different chromosomes (Supplementary Figure S11C). These correlations were especially pronounced when focusing on the top upregulated and downregulated genes/loci (Supplementary Figure S11D).

For gene markers from all 4 cell types in the adult mouse brain cortex (identified from RNA), we found that the corresponding scCompartments were more activated in their respective cell types, consistent with findings using scA/B (Supplementary Figure S12). Moreover, we computed the Spearman correlation between single-cell RNA expression and scCompartments. Compared to scA/B, our method demonstrated higher correlations across most chromosomes (Fig. 3D, Supplementary Figure S13).

To further validate this connection at single-cell resolution, we used an additional dataset, GAGE-seq on mouse brains [38]. Higher gene expression in a cell often corresponded to a higher real-valued compartment (Supplementary Fig-ure S13D). For 1,913 markers with significantly higher expressions in inhibitory neurons, most showed elevated compartmental values in these neurons compared to excitatory neurons (Supplementary Figure S13D). Thus, the relationship between compartment and gene expression remained evident at single-cell resolution.

We then illustrated these observations on a specific locus. Following the process in [38], we selected the gene exhibiting the most significant increase in compartmental value (by scDIAGRAM) and RNA expression in inhibitory neurons compared with excitatory neurons. This analysis led us to the same gene Erbb4 as in [38] when we looked at chr1 (Supplementary Figure S14A, B). In pseudo-bulk data, the Erbb4 locus switched from the stable B compartment (in excitatory neurons) to the stable A compartment (in inhibitory neurons). According to prior studies, the Erbb4 gene was essential in the central nervous system and has been associated with schizophrenia [45]. As expected, we observed differential A/B compartment values for excitatory and inhibitory neurons, correlated with cell-type-specific expression of the Erbb4 gene (Fig. 3E). The Pearson correlation of scCompartments and scRNAseq was higher inside inhibitory neurons (Fig. 3E). Additionally, we identified similar compartmental dynamics for the gene Npas3 on chr12 (Supplementary Figure S14C, D). This gene is a bHLH transcription factor regulating astrocyte-neuron communication and associated with autism [46]. It displayed a variable B to variable A compartmental switch from excitatory to inhibitory neurons, accompanied by increase in the transcriptional activity. Compared with Erbb4, this variable switch showed a lower Pearson correlation between scCompartments and scRNAseq.

### 3.3 scDIAGRAM applied to the developing mouse embryos

We applied scDIAGRAM to the HiRES dataset during embryogenesis.

#### 3.3.1 Compsc-Ncut of Ncut represents compartmental strength

The optimal 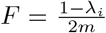 calculated from the second eigenvalue *λ*_*i*_ in normalized cut and the total edge weight *m* of the graph reflects how well the genome is partitioned into two compartments (See Supplementary Methods). We define −log_2_(*F*) as a quantitative measure of compartmental strength, denoted as Compsc-Ncut. Lower Compsc-Ncut values indicate weaker or less distinct compartmental separation.

We conducted permutation experiments to validate the ability of Compsc-Ncut from Ncut to represent compartmental strength of a Hi-C matrix. We randomly shuffled a Hi-C matrix and mixed it with the original one, breaking the compartmental structure. The Compsc-Ncut decreased as more shuffled matrices were mixed together (Supplementary methods and Supplementary Figure S15). In a real scHi-C dataset, we observed a very diversed range of the Compsc-Ncut (Fig. 4A), and we typically used a cutoff value of Compsc-Ncut at 17.3 in practical applications. We also presented two examples to illustrate the effect of Compsc-Ncut, where a lower Compsc-Ncut indicated a weaker compartmental structure (Fig. 4A).

**Figure 4:**
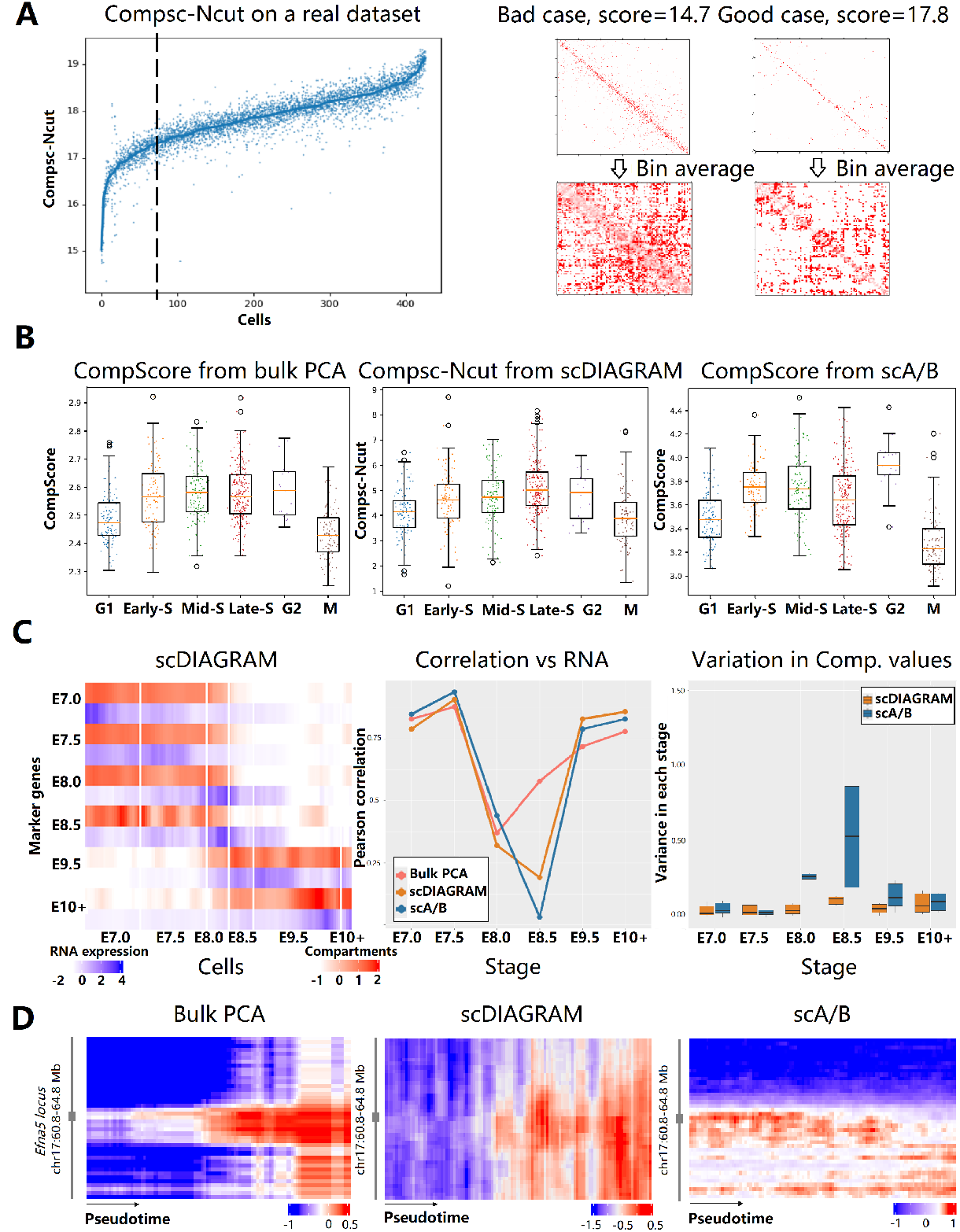
scDIAGRAM applied to the mouse embryonic development. (A). Compsc-Ncut of scDIAGRAM on a real scHi-C dataset and two examples, with larger values of Compsc-Ncut indicating clearer compartmental structure. Each cell was repeated for 10 times and the line was the averaged Compsc-Ncut for each cell. (B). Compartmental strength changes during cell cycle, using the Compsc-Ncut in scDIAGRAM and compartment score [11] for bulk PCA and scA/B. (C). Compartmental values from scDIAGRAM and RNA expressions (left), the Pearson correlation between compartmental values and RNA expressions (Middle), and the variance of compartmental values (Right) for each set of marker genes at different developmental stage. For each set of marker genes, the compartment annotations are averaged at each metacell. In the heatmap, each row represented a set of marker genes for a specific stage, while each column corresponded to a metacell, ordered according to pseudotime. (D) Compartments of the gene Efna5 and its neighboring loci at 100 kb resolution. Each column represents a cell, ordered by the pseudotime from left to right. Heatmaps were smoothed by every 5 neighboring metacells.

With this tool in hand, we were able to investigate the compartmental variation throughout the cell cycle. In Figure 4B, we used this Compsc-Ncut as a metric for compartmental strength and tracked its changes throughout the cell cycle. We observed a trend that closely mirrored the acknowledged compartmental change during the cell cycle, which was computed from the compartment score from bulk PCA [11]. In contrast, applying the compartment score to compartments generated by scA/B did not reveal the same trend, with only the maximum and minimum values aligning with what we expected.

#### 3.3.2 Dynamic compartmental evolution during development

The HiRES dataset included developing embryos spanning embryonic day 7.0 (E7.0) to day 11.5 (E11.5). scRNA-seq analysis revealed two developmental lineages: the neural and mesenchymal lineages, both originating from the epiblast and primitive streak [37]. To mitigate intrinsic noise and cell-cycle effects in the scHi-C data, we constructed “metacells” by aggregating single cells with similar expression profiles followed by trajectory inference and pseudotime analysis (Supplementary methods). In this study, we specifically focused on the neuronal trajectory to investigate compartmental transition dynamics during development.

Building on similar analyses from [38, 40], we examined compartmental changes associated with stage-specific marker genes. Given the continuous nature of embryogenesis and the potential ambiguity in cell type definitions, we identified marker genes separately for each embryonic stage. For each stage, all other stages were treated as controls, and differential expression analysis was performed using the Seurat package (see Supplementary methods). Subsequent analyses focused on the top 100 marker genes at each stage, ordered by the fold change.

In Fig. 4C, we visualized compartmental transitions of these marker genes during development, averaging the compartment values within each marker gene set. We assigned compartments for each metacell using bulk PCA, sc-DIAGRAM, and scA/B. For bulk PCA, compartments were assigned based on pseudo-bulk matrices constructed from cells at each developmental stage.

Compartmental changes exhibited a strong Pearson correlation with RNA expression for marker genes at the early (E7.0, E7.5) and late (E9.5, E10+) stages of development (Supplementary Figure S16). In contrast, markers at intermediate stages (E8.0–E8.5) showed markedly reduced correlations (Fig. 4C, middle). Marker genes specific to particular developmental stages generally displayed more active scCompartments, except at E8.5, where all methods produced highly variable scCompartments with only slight enrichment (Fig. 4C, right). These observations are consistent with the known intermixing of ectodermal and mesodermal cells prior to E8.5, which begin to diverge only afterward [37], further highlighting the cellular heterogeneity at this stage. Additionally, across developmental stages, scDIAGRAM showed better alignment with bulk PCA than scA/B (Supplementary Figure S16).

A closer examination of stage E8.0 revealed notable differences between sc-DIAGRAM and scA/B. We identified the gene Efna5, which showed opposing compartmental transitions when comparing bulk PCA and scA/B (Fig. 4D). In contrast to scA/B, scDIAGRAM aligned with bulk PCA, displaying an elevated compartmental signal, while scA/B indicated a more inactive state at this locus. Efna5 encodes an axon guidance protein involved in late-stage nervous system development by preventing axon bundling [47]. As shown in Fig. 4D, the compartment patterns of neighboring loci also more closely resembled those from bulk PCA when inferred by scDIAGRAM. scRNA-seq analysis further revealed that Efna5 expression increased at early developmental stages and declined thereafter, potentially explaining the discordance between expression and the compartment patterns detected by scA/B, as previously noted in [48].

### 3.4 scDIAGRAM applied to the acute myeloid leukaemia

We applied scDIAGRAM to the HiRES dataset of human AML. Previous bulk analyses had identified subtype-specific A/B compartment patterns in AML [49]. Consistent with this, scRNA-seq embeddings revealed patient-specific clustering (Supplementary Figure S17). We therefore focused our downstream analyses on compartmental differences across patients.

For chromosome 10 in Patient 03 (PT03) (Fig. 5A), scDIAGRAM demonstrated stronger concordance with bulk PCA than scA/B, as indicated by a higher Pearson correlation. Notably, in a region where scA/B showed substantial deviation from the bulk reference, scDIAGRAM remained closely aligned. In Fig. 5B (chromosome 10 across all patients), scDIAGRAM also exhibited greater compartmental heterogeneity compared to other methods. Following the approach in Fig. 3B, we computed the variance of scCompartments across cells. At the top 10% most variable loci—defined by bulk PCA—scDIAGRAM consistently showed higher variability than scHiCluster, scA/B, and Higashi across all patients.

**Figure 5:**
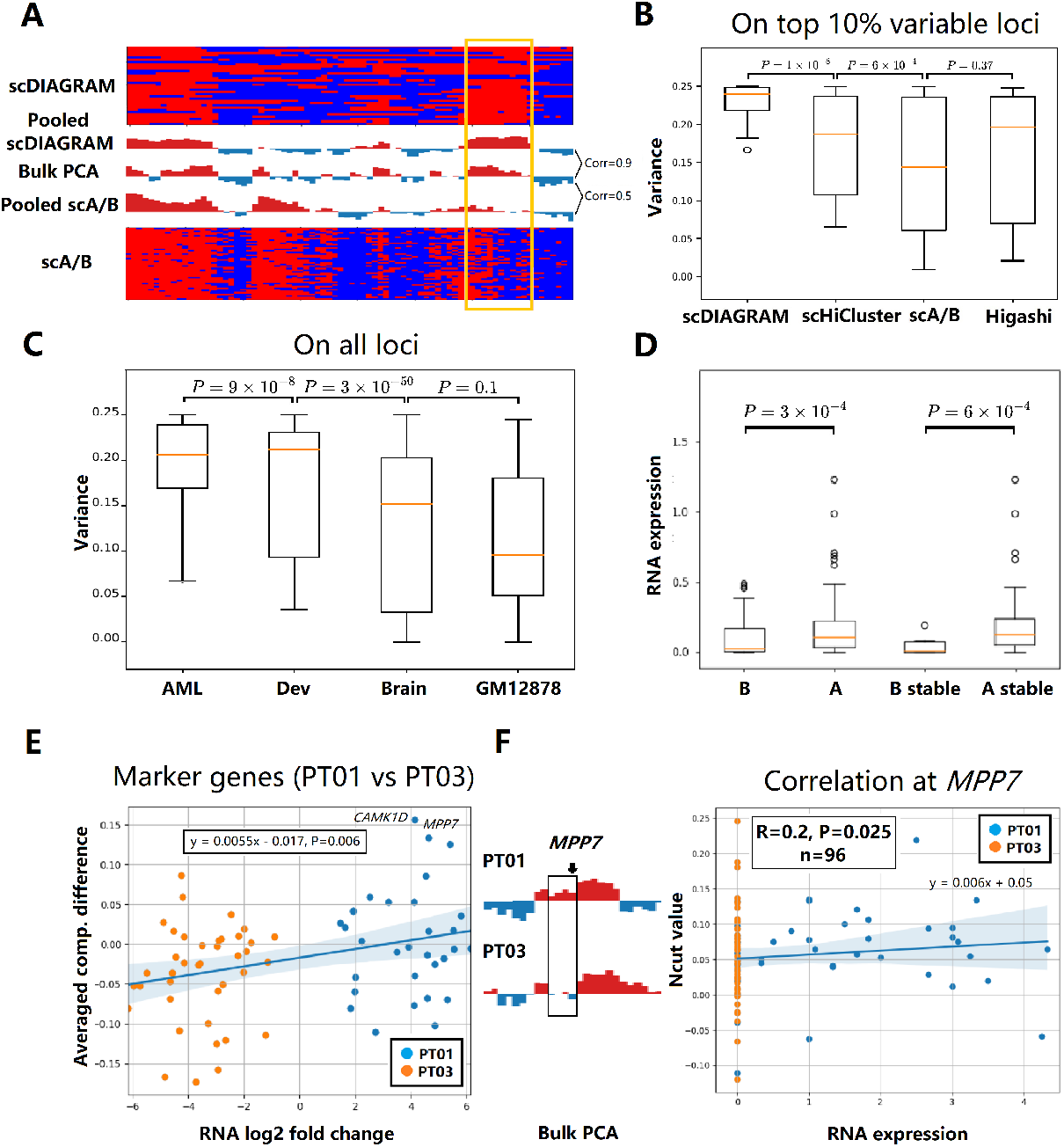
scDIAGRAM applied to the AML. (A) scCompartments inferred by scDIAGRAM showed closer agreement with bulk PCA than those from scA/B in the HiRES dataset of patient PT03. Each row represented a single cell. The highlighted region marked a locus where scA/B substantially deviated from bulk PCA. (B) scDIAGRAM captured greater compartmental heterogeneity compared to scHiCluster, Higashi, and scA/B. (C) Compartmental variance of scDIAGRAM was evaluated across different datasets. (D) Average RNA expression levels were compared between loci assigned to A and B compartments, as well as among stable A/B loci. (E) Changes in compartmentalization were correlated with gene expression differences for marker genes between patients PT01 and PT03. (F) Bulk PCA signals at the MPP7 locus were visualized for PT01 and PT03, along with corresponding RNA expression and scDIAGRAM Ncut values at the single-cell level.

Furthermore, cross-dataset comparisons using scDIAGRAM (Fig. 5C) revealed that the datasets of AML and developing embryo exhibited significantly higher scCompartment variability than both brain cells and the GM12878 cell line. This pattern is consistent with biological intuition: cancer cells and developing cells are expected to exhibit greater heterogeneity in chromatin organization due to their dynamic regulatory states, whereas terminally differentiated or steady-state cells tend to be more stable and homogeneous.

We next examined the relationship between compartments and transcriptional activity. As shown in Fig. 5D, loci annotated as A compartments were generally more transcriptionally active than B compartments. Moreover, stable A/B loci exhibited greater differences in gene expression, highlighting the functional relevance of compartmental identity. In Fig. 5E, comparing patients PT01 and PT03, the RNA fold change of all marker genes (*n* = 70, adjusted *p <* 0.05) was significantly correlated with the corresponding average compartmental differences (Spearman’s *r* = 0.32, *p* = 0.003). This association was consistently observed across other patient comparisons as well (Supplementary Figure S17).

Finally, in Fig. 5F, using the same approach as in Fig. 3E, we identified the gene MPP7, which exhibited the most pronounced increase in both compartmental value and RNA expression in PT01 compared to PT03 (Supplementary Figure S17). Bulk PCA analysis confirmed a B-to-A compartment switch at this locus between the two patients. This shift was mirrored by changes in MPP7 expression and scDIAGRAM-inferred compartment values, suggesting a coordinated regulatory transition. While MPP7 has been previously implicated in tumorigenesis, including in breast cancer [50], our findings suggest it may also play a role in AML, offering a novel biological prediction. A recent pan-cancer analysis based on public cancer databases has examined MPP7 across various cancer types, including AML [51], supporting the potential relevance of this gene in leukemogenesis. In addition to MPP7, we also identified known AML-associated genes, such as CAMK1D [52], that showed similar RNA and compartmental differences between PT01 and PT03 (Supplementary Figure S17).

## 4 Conclusion

In this study, we introduced scDIAGRAM, a computational method designed to annotate single-cell compartments directly from scHi-C data, without relying on CpG signals or imputation techniques. scDIAGRAM annotates compartments independently for each scHi-C matrix, without incorporating information from other cells. This approach enables us to capture the inherent heterogeneity of compartments within a scHi-C dataset, providing a more accurate representation of chromatin organization across diverse cell types. Through extensive simulations and real data analysis, we demonstrated that scDIAGRAM effectively annotated compartments across various cell types. We also identified cell-type-specific compartments and explored their association with gene expression profiles in the HiRES dataset.

Methods like Higashi [21] and scGHOST [40] have demonstrated the potential of artificial intelligence and deep neural networks (DNNs) for annotating compartments and subcompartments within scHi-C datasets. While DNN-based approaches generally require large datasets for training to achieve high performance by leveraging shared information across cells, the “black-box” nature of these models makes it challenging to interpret how compartment annotations are derived. In contrast, scDIAGRAM functions as a plug-and-play method, efficiently analyzing individual scHi-C matrices without the need for extensive training data. This makes scDIAGRAM an appealing option for users interested in studying compartmental heterogeneity and gene regulatory relationships, without the complexities associated with DNN-based methods.

The relationship between compartments, epigenomic features, and transcription is highly complex. Further quantitative analyses are needed to fully uncover these connections, and integrating epigenetic data with Hi-C could help improve the accuracy of compartment annotations. Additionally, understanding the mechanisms driving cell-to-cell compartmental variability is a significant challenge. This variability may stem from a range of factors, including cell type, cellular states, biological processes (such as the cell cycle), intrinsic cellular dynamics, and technical biases. Future research should focus on disentangling these factors and minimizing technical biases in scDIAGRAM, ultimately enhancing its robustness.

## Supporting information

Supplementary Methods, Results and Figures

## 5 Acknowledgments

We would like to thank Yue Xue, Bin Dong and Ruibin Xi for helpful discussions.

## 6 Author Contributions

H.G. conceived the project. Y.P. designed and implemented the scDIAGRAM algorithm under the supervision of H.G. and J.J.. Y.P. conducted the validation and simulations with assistance from Z.L. and D.X.. Y.D. and M.L. performed the HiRES experiments and data processing for the AML under supervision of D.X.. Y.L. and X.Z. collected AML samples. Y.P. and H.G. wrote the initial manuscript. All authors read and approved the final manuscript.

## 9 Data availability

The single-cell AML data generated in this study have been deposited in Gene Expression Omnibus (GEO) under accession code GSE302267. Other data are publicly available, and source code of scDIAGRAM are on Github (https://github.com/Ge-lab-pku/scDIAGRAM) and Zenodo (https://doi.org/10.5281/zenodo.15855256).

## References

[1] Erez Lieberman-Aiden, Nynke L. van Berkum, Louise Williams, and et al. Comprehensive mapping of long-range interactions reveals folding principles of the human genome. Science, 326(5950):289–293, 2009.

[2] Tom Misteli. The self-organizing genome: principles of genome architecture and function. Cell, 183:28–45, 2020.

[3] Zhijun Duan, Mirela Andronescu, Kevin Schutz, and et al. A three-dimensional model of the yeast genome. Nature, 465:363–367, 2010.

[4] Nicole Rusk. When ChIA-PETs meet Hi-C. Nat Methods, 6:863, 2009.

[5] M. J. Fullwood, M. Liu, Y. Pan, and et al. An oestrogen-receptor-α-bound human chromatin interactome. Nature, 462:58–64, 2009.

[6] T. Cremer and C. Cremer. Chromosome territories, nuclear architecture and gene regulation in mammalian cells. Nat. Rev. Genet., 2:292–301, 2001.

[7] Jesse R. Dixon, Siddarth Selvaraj, Feng Yue, and et al. Topological domains in mammalian genomes identified by analysis of chromatin interactions. Nature, 485:376–380, 2012.

[8] E. P. Nora, B. R. Lajoie, E. G. Schulz, and et al. Spatial partitioning of the regulatory landscape of the x-inactivation centre. Nature, 485:381–385, 2012.

[9] Suhas S.P. Rao, Miriam H. Huntley, Neva C. Durand, and et al. A 3D map of the human genome at kilobase resolution reveals principles of chromatin looping. Cell, 159:1665–1680, 2014.

[10] V. Ramani, X. Deng, R. Qiu, and et al. Massively multiplex single-cell Hi-C. Nat. Methods, 14:263–266, 2017.

[11] T. Nagano, Y. Lubling, C. Várnai, and et al. Cell-cycle dynamics of chromosomal organization at single-cell resolution. Nature, 547:61–67, 2017.

[12] Longzhi Tan, Dong Xing, Chi-Han Chang, Heng Li, and X. Sunney Xie. Three-dimensional genome structures of single diploid human cells. Science, 361:924–928, 2018.

[13] Hyeon-Jin Kim, Galip Gürkan Yardımcı, Giancarlo Bonora, and et al. Capturing cell type-specific chromatin compartment patterns by applying topic modeling to single-cell Hi-C data. PLoS Comput. Biol., 16:e1008173, 2020.

[14] D.-S. Lee, C. Luo, J. Zhou, and et al. Simultaneous profiling of 3D genome structure and dna methylation in single human cells. Nat. Methods., 16:999–1006, 2019.

[15] Takashi Nagano, Yaniv Lubling, Tim J Stevens, and et al. Single-cell Hi-C reveals cell-to-cell variability in chromosome structure. Nature, 502:59–64, 2013.

[16] Elizabeth H. Finn, Gianluca Pegoraro, Hugo B. Brandão, and et al. Extensive heterogeneity and intrinsic variation in spatial genome organization. Cell, 176:1502–1515.e10, 2019.

[17] Elizabeth H. Finn and Tom Misteli. Molecular basis and biological function of variability in spatial genome organization. Science, 365:eaaw9498, 2019.

[18] Tim J. Stevens, David Lando, Srinjan Basu, and et al. 3D structures of individual mammalian genomes studied by single-cell Hi-C. Nature, 544:59– 64, 2017.

[19] M. V. Arrastia, J. W. Jachowicz, N. Ollikainen, and et al. Single-cell measurement of higher-order 3D genome organization with scsprite. Nat. Biotechnol., 40:64–73, 2021.

[20] Jingtian Zhou, Jianzhu Ma, Yusi Chen, and et al. Robust single-cell Hi-C clustering by convolution-and random-walk–based imputation. Proc. Natl Acad. Sci., 116:14011–14018, 2019.

[21] Ruochi Zhang, Tianming Zhou, and Jian Ma. Multiscale and integrative single-cell Hi-C analysis with Higashi. Nat. Biotechnol., 40:254–261, 2022.

[22] Ruochi Zhang, Tianming Zhou, and Jian Ma. Ultrafast and interpretable single-cell 3D genome analysis with Fast-Higashi. Cell. Syst., 13:798–807, 2022.

[23] Yueying He, Yue Xue, Jingyao Wang, and et al. Diffusion-enhanced characterization of 3D chromatin structure reveals its linkage to gene regulatory networks and the interactome. Genome Res, 33:1354–1368, 2023.

[24] Ran Jiang, Yue Xue, Yanyi Huang, and Yiqin Gao. Deciphering single-cell 3D chromatin structure using scCTG. J. Chem. Phys., 161(24):245101, 2024.

[25] Yuxiang Zhan, Francesco Musella, and Frank Alber. Prediction of single-cell chromatin compartments from single-cell chromosome structures by maxcomp. bioRxiv, 2024.

[26] M. Yu, A. Abnousi, Y. Zhang, and et al. SnapHiC: a computational pipeline to identify chromatin loops from single-cell Hi-C data. Nat. Methods., 18:1056–1059, 2021.

[27] Y. Zhang, L. Boninsegna, M. Yang, and et al. Computational methods for analysing multiscale 3D genome organization. Nat. Rev. Genet., 25:123– 141, 2023.

[28] Tianming Zhou, Ruochi Zhang, and Jian Ma. The 3D genome structure of single cells. Annu. Rev. Biomed. Data Sci., 4:21–41, 2021.

[29] Angsheng Li, Guangjie Zeng, Haoyu Wang, and et al. DeDoc2 identifies and characterizes the hierarchy and dynamics of chromatin tad-like domains in the single cells. Advanced Science, 10(20):2300366, 2023.

[30] Siyuan Wang, Jun-Han Su, Brian J. Beliveau, and et al. Spatial organization of chromatin domains and compartments in single chromosomes. Science, 353:598–602, 2016.

[31] Yodai T., Yujing Yang, Jonathan White, and et al. High-resolution spatial multi-omics reveals cell-type specific nuclear compartments. biorxiv, 2023.

[32] Open2C, Nezar Abdennur, Sameer Abraham, and et al. Cooltools: Enabling high-resolution Hi-C analysis in python. PLOS Computational Biology, 20(5):1–16, 05 2024.

[33] M. Simonis, P. Klous, E. Splinter, and et al. Nuclear organization of active and inactive chromatin domains uncovered by chromosome conformation capture-on-chip (4c). Nat. Genet., 38:1348–1354, 2006.

[34] Wen Jun Xie, Luming Meng, Sirui Liu, and et al. Structural modeling of chromatin integrates genome features and reveals chromosome folding principle. Sci Rep, 7:2818, 2017.

[35] M. E. J. Newman. Spectral methods for community detection and graph partitioning. Phys. Rev. E, 88:042822, 2013.

[36] Zhihua Zhang and Michael I. Jordan. Multiway spectral clustering: A margin-based perspective. Stat. Sci., 23(3):383–403, 2008.

[37] Zhiyuan Liu, Yujie Chen, Qimin Xia, and et al. Linking genome structures to functions by simultaneous single-cell Hi-C and RNA-seq. Science, 380:1070–1076, 2023.

[38] Tianming Zhou, Ruochi Zhang, Deyong Jia, and et al. GAGE-seq concurrently profiles multiscale 3D genome organization and gene expression in single cells. Nat. Genet., 56:1701–1711, 2024.

[39] Y. Hao, S. Hao, E. Andersen-Nissen, and et al. Integrated analysis of multimodal single-cell data. Cell, 184:3573–3587.e29, 2021.

[40] K. Xiong, R. Zhang, and J. Ma. scGHOST: identifying single-cell 3D genome subcompartments. Nat Methods, 21:814–822, 2024.

[41] Jun-Han Su, Pu Zheng, Seon S. Kinrot, and et al. Genome-scale imaging of the 3D organization and transcriptional activity of chromatin. Cell, 182:1641–1659, 2020.

[42] L. Chang, Y. Xie, B. Taylor, and et al. Droplet Hi-C enables scalable, single-cell profiling of chromatin architecture in heterogeneous tissues. Nat Biotechnol, 2024.

[43] P. Trojer and D. Reinberg. Facultative heterochromatin: is there a distinctive molecular signature? Mol. Cell, 28:1–13, 2007.

[44] Chenxu Zhu, Yanxiao Zhang, Yang Eric Li, and et al. Joint profiling of histone modifications and transcriptome in single cells from mouse brain. Nat. Methods., 18:283–292, 2021.

[45] Amanda J Law, Joel E Kleinman, Daniel R Weinberger, and Cynthia Shannon Weickert. Disease-associated intronic variants in the ErbB4 gene are related to altered ErbB4 splice-variant expression in the brain in schizophrenia. Hum. Mol. Genet., 16:129–141, 2007.

[46] Yuanyuan Li, Tianda Fan, Xianfeng Li, and et al. Npas3 deficiency impairs cortical astrogenesis and induces autistic-like behaviors. Cell Reports, 40:111289, 2022.

[47] John W. Winslow, Paul Moran, Janet Valverde, and et al. Cloning of AL1, a ligand for an Eph-related tyrosine kinase receptor involved in axon bundle formation. Neuron, 14(5):973–981, 1995.

[48] Longzhi Tan, Wenping Ma, Honggui Wu, and et al. Changes in genome architecture and transcriptional dynamics progress independently of sensory experience during post-natal brain development. Cell, 184:741–758.e17, 2021.

[49] J. Xu, F. Song, H. Lyu, and et al. Subtype-specific 3D genome alteration in acute myeloid leukaemia. Nature, 611:387–398, 2022.

[50] W. Liao, L. Fan, M. Li, and et al. MPP7 promotes the migration and invasion of breast cancer cells via EGFR/AKT signaling. Cell Biology International, 45(5):948–956, 2021.

[51] X. Xu, W. Cheng, S. Zhao, and et al. Pan-cancer analysis of the role of MPP7 in human tumors. Heliyon, 10(16):e36148, 2024.

[52] X. Kang, C. Cui, C. Wang, and et al. CAMKs support development of acute myeloid leukemia. J Hematol Oncol, 11:30, 2018.

