## Supplementary Methods, Results and Figures for "scDIAGRAM: Detecting Chromatin Compartments from Individual Single-Cell Hi-C Matrix without Imputation or Reference Features"

#### 1 Supplementary Methods

##### 1.1 The derivation and details of normalized cut

The objective function can be expressed into a quadratic form:

$$\frac{R}{\kappa_1\kappa_2} = \frac{2m - \mathbf{s}^T\mathbf{A}\mathbf{s}}{(2m)^2}, \quad s_i = \begin{cases} \sqrt{\kappa_2/\kappa_1} & \text{if} \quad i \text{ is in compartment } 1, \\ -\sqrt{\kappa_1/\kappa_2} & \text{if} \quad i \text{ is in compartment } 2, \end{cases}$$

where $\mathbf{A}$ is the adjacency matrix and $m$ represents the total weight of all edges in the graph.

Next, we relaxed the above discrete $\mathbf{s}$ into a vector in $\mathbb{R}^n$, transforming the problem into:

$$\max_{\mathbf{s}\in\mathbb{R}^n} \mathbf{s}^T\mathbf{A}\mathbf{s}, \text{ s.t. } \sum_i k_i s_i = 0, \sum_i k_i s_i^2 = 2m,$$

where $k_i$ is the degree of node $i$.

This problem can be solved by introducing Lagrange multipliers $\lambda, \mu$ for the two constraints and differentiating, i.e.

$$\mathbf{A}\mathbf{s} = \lambda\mathbf{D}\mathbf{s} + \mu\mathbf{k},$$

where $\mathbf{k}$ is the vector with elements $k_i$ and $\mathbf{D}$ is the diagonal matrix with elements $D_{ii} = k_i$. Multiplying on the left by $\mathbf{1}^T$ and using $\mathbf{1}^T\mathbf{A} = \mathbf{1}^T\mathbf{D} = \mathbf{k}^T$, $\mathbf{k}^T\mathbf{s} = 0$ one got $\mu = 0$. Hence the solution should satisfy the generalized eigenvector equation

$$\mathbf{A}\mathbf{s} = \lambda\mathbf{D}\mathbf{s}.$$

This equation can be explicitly solved by considering the eigen decomposition of $\mathbf{D}^{-1/2}\mathbf{A}\mathbf{D}^{-1/2}$. We denoted the solutions to this generalized eigenvector equation were $(\nu_1, \nu_2, \cdots, \nu_{K+1})$ with the corresponding eigenvalues $(\lambda_1 \geq \lambda_2 \geq \cdots \geq \lambda_{K+1})$. The following properties hold:

- each $\lambda_i \leq 1$, $\lambda_1 = 1$ and $\nu_1 = \mathbf{1} = (1, 1, 1 \cdots)^T$;
- $\frac{R}{\kappa_1\kappa_2} = \frac{2m - \mathbf{s}^T\mathbf{A}\mathbf{s}}{(2m)^2} = \frac{1-\lambda_i}{2m}$ if $\mathbf{s} = \nu_i$ and $\sum_i k_i s_i^2 = \mathbf{s}^T\mathbf{D}\mathbf{s} = 2m$, for any $i$.

Since $\nu_1 = \mathbf{1}$ can not satisfy the constraint $\mathbf{k}^T\mathbf{s} = 0$, we turned to $\nu_2$, which is exactly the solution of the optimization problem.

To obtain a discrete $\mathbf{s}$ for graph partitioning, $\mathbf{s}$ was rounded, typically using 0 as a threshold and nodes were assigned to the two compartments based on the sign of $\mathbf{s}$ (according to Newman (2013), the result was robust to the rounding strategy chosen for division). The compartmental value of each locus just inherited from the value of the corresponding node, i.e. the group of loci separated by CPs.

##### 1.2 MCMC algorithm for the posterior distribution

The pseudocode was provided in Algorithm 1.

To be specific, we used $v_i = 1$ to denote the presence of a CP between the $i$-th and $(i+1)$-th loci, and $v_i = 0$ otherwise. Starting from the current state $(v_i)_{1\leq i\leq n-1}$, we randomly selected $j \in I_1 = \{i : v_i = 1\}$, $k \in I_0 = \{i : v_i = 0\}$ and proposed the next candidate state $(v_i')_{1\leq i\leq n-1}$ which moves one CP at position $j$ to position $k$. Each pair $(j, k)$ had equal probability $\frac{1}{2K(n-1-K)}$ with $|I_1| = K$. This proposal was symmetric.

We accepted this candidate $(v_i')_{1\leq i\leq n-1}$ with probability $\alpha(v_1, \cdots, v_{n-1}; v_1', \cdots, v_{n-1}')$ (defined in Supplementary Methods 1.3), which was derived from the likelihood ratio between the two states. In

**Algorithm 1:** Metropolis-Hasting Sampling

**Input:** Matrix $\{Y_{ij}\}$, number of iterations $N$, number of changing points $K$;

Initialize the positions of all CPs $(v_i^{(0)})$ (randomly choose K positions as CPs), $t = 0$;

**Output:** $(v_i^{(1)}), \cdots, (v_i^{(N)})$;

**for** $t = 0, 1, \ldots, N-1$ **do**

  Randomly select $j$ from $I_1 = \{i : v_i^{(t)} = 1\}$ and $k$ from $I_0 = \{i : v_i^{(t)} = 0\}$;

  define $v_i' = \begin{cases} 1 - v_i^{(t)}, & \text{if } i = j \text{ or } k \\ v_i^{(t)}, & \text{otherwise} \end{cases}$

  $p_{acc} = \alpha(v_1^{(t)}, \cdots, v_{n-1}^{(t)}; v_1', \cdots, v_{n-1}')$ # $\alpha$ defined in Appendix;

  **if** $u \sim U(0,1) \leq p_{acc}$ **then**

    Accept candidate $(v_i^{(t+1)}) \leftarrow (v_i')$

  **else**

    Reject candidate $(v_i^{(t+1)}) \leftarrow (v_i^{(t)})$

  **end**

**end**

this way, the sampling sequence formed a reversible Markov chain and the distribution converged to the posterior distribution in the limit.

Since MCMC involves inherent randomness, we repeated the process multiple times (typically 5–10 repetitions) to improve robustness.

Finally, we selected the state with the maximal likelihood value among all the sampled states as an approximation to the MLE solution.

#### 1.3 The acceptance probability in MCMC

Specifically, given the present state $(v_i)_{1 \leq i \leq n-1}$, we randomly moved one CP at the position $j$ to an unoccupied position $k$, proposing a candidate state.

We accepted this proposal $(v_i')_{1 \leq i \leq n-1}$ with probability $\alpha(v_1, \cdots, v_{n-1}; v_1', \cdots, v_{n-1}')$, which was the minimum of the likelihood ratio between these two states and 1.

The likelihood was just the posterior (defined in Methods of the main text), so

$$\begin{aligned} &\alpha(v_1, \cdots, v_{n-1}; v_1', \cdots, v_{n-1}') \\ = \quad &\min\left\{1, \frac{\prod_{l \neq k, k+1} A(v_i, k, l)}{\prod_{l \neq j, j+1} A(v_i, j, l)} \times \frac{B(v_i, k)}{B(v_i, j)}\right\}, \end{aligned} \quad (1)$$

in which

$$\begin{aligned} A(v_i, k, l) = &\frac{\hat{r}_{kl}^{S_{kl}} \hat{r}_{k+1,l}^{S_{k+1,l}}}{(\alpha_{kl}\hat{r}_{kl} + (1-\alpha_{kl})\hat{r}_{k+1,l})^{S_{kl}+S_{k+1,l}}} \\ \times &\frac{(1-\hat{r}_{kl})^{N_{kl}-S_{kl}}(1-\hat{r}_{k+1,l})^{N_{k+1,l}-S_{k+1,l}}}{[\alpha_{kl}(1-\hat{r}_{kl}) + (1-\alpha_{kl})(1-\hat{r}_{k+1,l})]^{N_{kl}+N_{k+1,l}-S_{kl}-S_{k+1,l}}} \\ \times &\frac{\exp\left(-\frac{1}{2\hat{\sigma}^2}(S_{kl}m_{kl}^2 + S_{k+1,l}m_{k+1,l}^2)\right)}{\exp\left(-\frac{1}{2\hat{\sigma}^2}(S_{kl} + S_{k+1,l})(\beta_{kl}m_{kl} + (1-\beta_{kl})m_{k+1,l})^2\right)}, \end{aligned}$$

where $\alpha_{kl} = \frac{N_{kl}}{N_{kl}+N_{k+1,l}}$, $\beta_{kl} = \frac{S_{kl}}{S_{kl}+S_{k+1,l}}$ and $m_{kl}$ was the sample mean of nonzero entries in the $(k,l)$-block. Defined as before, $N_{kl}$ was the sample size of the $(k,l)$-block and $S_{kl} = \sum_{i \in g_k, j \in g_l} 1_{\{X_{ij} \neq 0\}}$ was the nonzero sample size in the $(k,l)$-block.

The expression of $B(v_i, k)$ is

$$
\begin{aligned}
B(v_i,k) &= \left(\frac{\hat{r}_{kk}}{\overline{r}_k}\right)^{S_{kk}} \left(\frac{1-\hat{r}_{kk}}{1-\overline{r}_k}\right)^{N_{kk}-S_{kk}} \\
&\times \left(\frac{\hat{r}_{k+1,k+1}}{\overline{r}_k}\right)^{S_{k+1,k+1}} \left(\frac{1-\hat{r}_{k+1,k+1}}{1-\overline{r}_k}\right)^{N_{k+1,k+1}-S_{k+1,k+1}} \\
&\times \left(\frac{\hat{r}_{kk}}{\overline{r}_k}\right)^{2S_{k,k+1}} \left(\frac{1-\hat{r}_{k,k+1}}{1-\overline{r}_k}\right)^{2N_{k,k+1}-2S_{k,k+1}} \\
&\times \frac{\exp\left(-\frac{1}{2\hat{\sigma}^2}(S_{kk}m_{kk}^2 + S_{k+1,k+1}m_{k+1,k+1}^2 + 2S_{k,k+1}m_{k,k+1}^2)\right)}{\exp\left(-\frac{1}{2\hat{\sigma}^2}(S_{kk} + S_{k+1,k+1} + 2S_{k,k+1})\overline{m}_k^2\right)},
\end{aligned}
$$

where $\overline{r}_k = A_k\hat{r}_{kk} + B_k\hat{r}_{k+1,k+1} + 2C_k\hat{r}_{k,k+1}$ was the weighted average of $(\hat{r}_{kk}, \hat{r}_{k,k+1}, \hat{r}_{k+1,k+1})$, and $\overline{m}_k = D_k m_{kk} + E_k m_{k+1,k+1} + 2F_k m_{k,k+1}$ was the weighted average of $(m_{kk}, m_{k,k+1}, m_{k+1,k+1})$.

The weights $A_k, B_k, C_k, D_k, E_k, F_k$ were given by

$$
\begin{aligned}
A_k &= \frac{N_{k,k}}{N_{k,k} + N_{k+1,k+1} + 2N_{k,k+1}}; \\
B_k &= \frac{N_{k+1,k+1}}{N_{k,k} + N_{k+1,k+1} + 2N_{k,k+1}}; \\
C_k &= \frac{N_{k,k+1}}{N_{k,k} + N_{k+1,k+1} + 2N_{k,k+1}}; \\
D_k &= \frac{S_{k,k}}{S_{k,k} + S_{k+1,k+1} + 2S_{k,k+1}}; \\
E_k &= \frac{S_{k+1,k+1}}{S_{k,k} + S_{k+1,k+1} + 2S_{k,k+1}}; \\
F_k &= \frac{S_{k,k+1}}{S_{k,k} + S_{k+1,k+1} + 2S_{k,k+1}}.
\end{aligned}
$$

#### 1.4 Quality control of compartmental structure using scDIAGRAM

In some cells, the compartmental structure appeared ambiguous, making it necessary to exclude them from downstream analyses. We computed Compsc-Ncut for each cell and applied a cutoff (typically 17.0–17.3) to filter out cells with poorly defined compartment patterns.

We also used the Pearson correlation between the CpG density and the annotation generated by scDIAGRAM as an additional filtering criterion. A lower correlation value indicated a significant deviation from expected patterns. A threshold between 0.1 and 0.2 was typically sufficient to identify and exclude such cases.

#### 1.5 Compartmental strength and normalized cut

We compared the relation between the Compsc-Ncut and the original CompScore defined in Nagano et al. (2017). The original CompScore was:

$$
\text{CompScore} = \log_2\left(\frac{2\,(O_{AA}+O_{AB})\,(O_{BB}+O_{AB}))}{mO_{AB}}\right),
$$

where $O_{AA}, O_{AB}, O_{BB}$ were total contact numbers between A-A, A-B and B-B compartments, m was the total contacts in the scHi-C matrix (the edge number in the graph). From Newman (2013) we defined the Compsc-Ncut as:

$$
\begin{aligned}
\text{Compsc-Ncut} &= -\log_2(F), \\
F = \frac{O_{AB}}{\kappa_A\kappa_B} &= \frac{O_{AB}}{(O_{AA}+O_{AB})\,(O_{BB}+O_{AB})},
\end{aligned}
$$

where $\kappa_A, \kappa_B$ were the total contact number (the total degrees) in A/B compartment.
Therefore we can derive CompScore − Compsc-Ncut $= \log_2(2/m)$. When comparing compartment

strength in real scHi-C matrices or balanced bulk Hi-C matrices, if the total contacts didn't change much during the comparison, these two metrics were equivalent.

We implemented some permutation experiments to validate the relation between Compsc-Ncut and the compartmental strength. These permutations would disrupt the compartmental structure.

First, we performed random shuffling (reordering) of the genomic bins in the Hi-C matrix. Shuffling did not affect the graphical structure and the compartments were unchanged, but the binary division changed. To disrupt the compartmental structure, we then mixed the original data with the shuffled data (just adding them together). By adjusting the weight of the shuffled data upon mixing, we can also control the extent to which the compartmental structure was disrupted.

#### 1.6 Simulated scHi-C datasets

##### Downsampling of pseudo-bulk Hi-C data

To generate simulated scHi-C datasets, we performed downsampling on pseudo-bulk Hi-C data derived from the HiRES Liu et al. (2023) dataset, i.e. the excitatory neuron cluster 1 (Ex1) and mixed late mesenchyme (MLM) cell types at 100 kb resolution. The results from bulk PCA served as the ground truth of compartmentalization.

At a given downsampling rate 1/r, we randomly selected 1/r of all the sequencing reads in the pseudo-bulk matrix to create a sampled contact map. This approach was equivalent to applying binomial sampling. The downsampling procedure was repeated 50 times at each rate. The choice of downsampling rates was guided by the contact numbers observed in the pseudo-bulk data and real single-cell data. The rates used in this study were 1/400, 1/800, 1/1200 (close to real scHi-C dataset), 1/1600, 1/2400 and 1/3200.

##### Downsampling from single-cell 3D genome imaging data

We further simulated an additional scHi-C dataset using a recent 3D genome imaging dataset comprising 3,029 chromosomes (chr2) of single cells from Su et al. (2020). This dataset provided chromosome labeling at 250 kb resolution. We generated synthetic scHi-C matrices at 1 Mb resolution from this data, just like the processing steps in Zhang et al. (2022).

First we averaged the coordinates of every four 250 kb segments to derive the spatial coordinates of each 1 Mb segment. As demonstrated in Bintu et al. (2018), the inverse of the spatial distance was strongly correlated with Hi-C contact frequencies between pairs of genomic loci. Using this relationship, we constructed the original dataset by taking the inverse of the distance matrix (the diagonal all set to zero in the original data). The PCA of the original matrix was used as the ground truth for subsequent analysis.

Reads were then randomly sampled with probabilities proportional to the values in this ground-truth matrix. The law of large numbers ensured that the sampled contact matrice converged to the ground truth as the total contact number $n$ increased. We generated samples with contact numbers of 250, 500, 1,000, 2,500, 5,000, and 10,000. Supplementary Figure S4 listed the contact numbers per cell for various scHi-C datasets on the same chromosome. A typical scHi-C dataset had median contact number ranging from 500 to 5,000 while recent high coverage datasets had median contact number larger than 10,000.

##### Downsampling of high coverage scHi-C data

We further assessed the robustness of scDIAGRAM to sequencing depth by downsampling a high-coverage scHi-C dataset. We used the Dip-C dataset, which includes data from 14 GM12878 cells (chr1) at 500 kb resolution Tan et al. (2018). The corresponding 3D structural models were obtained using the Hickit package. Similar to the above approach for downsampling single-cell 3D imaging data, we took the inverse of the distance matrix and applied PCA to obtain the ground truth for this analysis. Following Zhang et al. (2022), 500 kb were chosen here so that Hickit can produce 3D structures at full coverage, as 3D structures at higher resolutions would skip bins with insufficient Hi-C contacts.

Next, we downsampled the Dip-C dataset to 50%, 25%, 10%, 5%, and 1% of the original read coverage, following the same procedure used for downsampling pseudo-bulk Hi-C data. The 1% read coverage corresponds to the median number of contacts per cell typically observed in scHi-C datasets (Supplementary Figure S4).

#### 1.7 Preprocessing of real scHi-C and scRNA-seq data

The mouse reference genome (GRCm38) and gene annotations (ALL) were downloaded from the GENCODE M23 release. The human reference genome (GRCh37 for GM12878 and GRCh38 for AML) and gene annotations were downloaded from UCSC Genome Browser Kent W.J and et al (2002). The CpG density data were computed from these reference genomes.

We used cooler Abdennur and Mirny (2020) and cooltools Open2C et al. (2024) to apply band normalization, matrix balancing, and compartment annotation to our pseudo-bulk Hi-C matrices. Single-cell Hi-C matrices also underwent band normalization prior to bin averaging. However, the extreme sparsity of raw scHi-C data makes it impossible to perform reliable band normalization or matrix balancing using only individual single-cell matrices. To overcome this, we recommend borrowing the expected contact frequency vectors and balancing weights calculated from the pseudo-bulk matrices when normalizing and balancing each single-cell dataset.

The HiRES dataset from Liu et al. (2023) was downloaded from the GEO (GSE223917). The data processing method for AML in this study follows the same procedure as in Liu et al. (2023). The GAGE-seq dataset was downloaded from the GEO (GSE238001). The imaging dataset Su et al. (2020) was obtained from Zenodo (https://doi.org/10.5281/zenodo.3928890). The Dip-C high coverage scHi-C dataset Tan et al. (2018) was downloaded from the GEO (GSE117876). The processed scHi-C data of the GM12878 cell line were downloaded from Kim et al. (2020) (https://noble.gs.washington.edu/proj/sc hic-topic-model), at 500 kb resolution. We transformed this dataset into 1 Mb resolution in our analysis. The epigenetic data of GM12878 were downloaded from ENCODE datasets: ENCFF167NBF (H3K27me3), ENCFF171MDW (H3K36me3), ENCFF803DJF (H3K79me2), ENCFF77 6OVW (H3K9me3) and ENCFF180LKW (H3K27ac).

##### HiRES

After the quality control filtering in HiRES, a total of 399 cells from mouse brain and 7,469 cells from mouse embryos were obtained. All mouse brain cells remained were used for analysis, while for mouse embryos we focused on the two primary lineages comprising of 3,247 cells. The cell types were provided in the dataset, obtained from annotations on scRNA-seq. For the AML dataset, a total of 427 cells were obtained from 6 patients (named as PT01-PT06).

The scHi-C matrices were binned at 100 kb in mouse brains and embryos; at 500 kb in AML, unless otherwise stated. Only autochromosomes (chr1-19 for mouse; chr1-22 for human) were used for analysis. We ran scDIAGRAM on each chromosome separately. The preprocessing of pseudo-bulk Hi-C matrices in scDIAGRAM precisely followed the standardized pipeline implemented in cooltools Open2C et al. (2024).

We constructed metacells on two lineages and computed pseudotime on the early neuronal lineage (consisting of 2,296 cells). All cells in mitosis (M stage) were excluded when constructing metacells. The marker genes in mouse brains, embryos and AML were identified using Seurat Hao et al. (2021) with default parameters. Details can be referred to Supplementary Methods 1.9.

##### GAGE-seq

The quality control filtering of the GAGE-seq followed the same thresholds in Zhou et al. (2024). We only used the mouse brain cortex dataset in GAGE-seq. Cells were retained if it had: 1) at least 1,000 mouse RNA reads, 2) at most 1% of RNA coming from mouse mitochondria, 3) at least 50,000 mouse contact pairs, 4) at least 20 contact pair per 1 Mb on average on each mouse chromosome. The scHi-C matrices were also binned at 100 kb and autochromosomes (chr1-19) were used for analysis.

GAGE-seq may generate doublets. We used the DoubletDetect tool Gayoso et al. (2022) to detect and remove doublets. The BoostClassifier was trained with parameters n_iters=100, n_components=28. Doublets were then inferred by the trained classifier with thresholds p_thr=1e-2, v_thr=.3. After this, cells with more than 45K nonzero elements in the contact map were removed.

The cell types of GAGE-seq were obtained from clustering and annotation of scRNA-seq from GAGE-seq. The marker genes were also identified using Seurat Hao et al. (2021) with default parameters. Details can be referred to Supplementary Methods 1.9.

##### scRNA-seq

When computing the coefficient of variation (CV) to measure the transcriptional variability.

In scRNA-seq data, there is an inherent connection between the mean and standard deviation of read counts Hafemeister and Satija (2019). Thus we used the coefficient of variation (CV, the ratio of the standard deviation to the mean), to measure the transcriptional variability.

Due to the dropout events in the scRNA-seq, we utilized MAGIC Van Dijk and et al. (2018) to first impute the scRNA-seq data. Later when computing the correlation of compartments and RNA we also implemented the same imputation (Fig 3D, Supplementary Figure S13). In other cases, the original scRNA-seq was used.

#### 1.8 Experimental details for the HiRES on AML

##### Patients source

We collected acute myeloid leukemia (AML) patient samples and their pathological results from the Department of Hematology at Peking University People's Hospital. All samples were obtained via bone marrow aspiration. The study was approved by the Ethics Committee of Peking University People's Hospital (2024PHB391-001). All patients signed informed consent forms as required.

##### Isolation of single cells

We isolated mononuclear cells from each patient's whole blood sample by using the Ficoll product (TBD, LTS1077). After density gradient centrifugation performed at 2000 × g for 20 minutes at 9:0 (brake off), cells were resuspended with 10 mL PBS (Gibco, 2124859). Then, we use 5 mL red blood cell lysis solution (Solarbio, 2312012) to remove red blood cells for 6 to 8 minutes on ice. The cells were resuspended with PBS and stained with flow cytometry antibodies CD34, CD117, CD45, 7-AAD, CD38 (Biolegend, 562577, 313218, 560777, 420403, 356605). After 20 minutes, cells were sorted by BD FACS Aria SORP to get CD34+CD117+CD38- hematopoietic stem and progenitor cells (HSPCs) for downstream HiRES experiments.

##### Library preparation and sequencing

Library was prepared strictly following HiRES protocolLiu et al. (2023). Sorted single cells were fixed with final 1.75% paraformaldehyde (PFA, ThermoFisher 28906) for 10 minutes at room temperature, followed by quenching with 2% BSA at a 10:1 volume ratio. Then, cells were resuspended in 200 $\mu$L ice-cold Wash Buffer (10 mM Tris pH 8.0, 10 mM NaCl, 0.1 mg/mL BSA (NEB B9000S)) supplemented with 20 $\mu$L protease inhibitor (Sigma P8340) and 2.6 $\mu$L Recombinant RNase Inhibitor. Each barcoded single-cell libraries were pooled and purified with 0.6× and 0.15× AMPure XP beads. The final libraries were sequenced with paired-end 150-bp reads on a NovaSeq X Plus (Illumina) platform.

#### 1.9 More details in Data processing

##### Cell type annotating in GAGE-seq

In the GAGE-seq dataset from mouse brains, we used its scRNA-seq for cell type annotating. This dataset were generated from cells in the mouse cortex (8–9 weeks old), consisting of 3,143 cells after filtering.

We used the Seurat Hao et al. (2021) package. First we selected genes expressed in at least 10 cells. The expression data were then normalized using the 'NormalizeData' and 'ScaleData' functions with default parameters. Highly variable genes were identified with the 'FindVariableFeatures' function. PCA analysis was performed with the 'RunPCA' function with npcs=50 PCs. The Louvain clustering was performed with functions 'FindNeighbors' and 'FindClusters' using 20 neighbors, the first 27 PCs, the euclidean distance as the metric, and a resolution of 3. Then we run UMAP with 'RunUMAP' on the first 25 PCs to generate cell embedding and clustering.

After clustering we generated 29 clusters. Using the marker genes from the original paper Zhou et al. (2024) (genes *Slc17a7*, *Gad1*, *Slc7a10*, *Cspg4*, *Mag*, *Apod*, *Cx3cr1*), we divided these 29 clusters into three major lineages in the mouse cortex: 16 excitatory neuron subtypes, 8 inhibitory neuron subtypes and 5 glial cell subtypes. Each lineage exhibited unique marker gene expressions. In later studies we would compare the transcription (RNA) and genome organization (Hi-C) between the excitotory and inhibitory lineage.

##### Detecting markers from scRNA-seq

In the HiRES data from mouse brains, we have cell type annotation provided by the authors, with 7 cell types in total (excitatory neuron cluster 1-3, Ex1-3; inhibitory neuron cluster 1-2, In1-2; astrocyte, Ast; oligodendrocyte, Oli). Since the compartmental enrichments were not that obvious and quite noisy between cell subtypes (both for scDIAGRAM and scA/B), we here focused on 4 major cell types: Ex, In, Ast and Oli. We just utilized these cell types and used the Seurat ”FindAllMarkers” function with ”only.pos=T” parameter. Thus for each cell type, we used the remaining cell types as control and only detected those up-regulated markers in this cell type. All the other parameters were the default parameters. In Supplementary Figure S12, we selected the top 500 markers for each cell type and computed their averaged scCompartments for each method.

In the GAGE-seq from mouse brains, after cell embedding and cell type annotating of the scRNA-seq, we only focused on the two large groups of cells, consisting of excitatory and inhibitory neurons. So we just used the Seurat ”FindMarker” function to detect up-regulated markers in inhibitory neurons. Here we used the parameters “p_val_adj $< 0.05$” and “min.pct $= 0.01$”. In this dataset we used the “MAST” test instead of the default “wilcox” test. All the other parameters were the default parameters.

In the HiRES data from the developing mouse embryos, we detected stage-specific markers for each stage, overlooking their cell types. We still used the the Seurat ”FindAllMarkers” function with ”only.pos=T” parameter, but we used the stages to group cells. So for each stage, we used the remaining stages as control and detected up-regulated markers for this stage. All the other parameters were the default parameters.

##### Embedding using scHi-C data

These embeddings were only used in Supplementary Figure S7A for UMAP visualization.

In the HiRES data from adult mouse brains, we compared different methods of cell embeddings based on scHi-C data (Supplementary Figure S7A).

First we used scCompartments from scDIAGRAM and scCpG for cell embedding. For scDIAGRAM we used the real-valued Ncut. First we computed PCA and selected the first 15 PCs. Then we run UMAP on these PCs, with the parameter “n_neighbors=10” and “min_dist=0”. All the other parameters were set as default.

For scCpG we first performed a rank normalization into [0,1] after concatenating results from chr1-19. Then we did the same things as above. Took the first 15 PCs from PCA and fed them into UMAP with exactly the same parameters.

Then we used scHiCluster imputed matrix for cell embedding. For scHiCluster, we just utilized the flattened imputed matrices for each chromosome, and only considered contacts between pairs of loci located within 10Mb on the genome. We took the first 50 PCs for each intrachromsomal Hi-C matrix, then we concatenated them and took the first 20 PCs again and fed them into UMAP with the same parameters.

Higashi automatically generated embeddings. We just used its embeddings and fed into UMAP with the same parameters.

##### Metacell construction

In the HiRES data from developing mouse embryos, to decrease noise in single-cell data, we generated metacells for downstream analysis, following the same steps in Liu et al. (2023).

We defined single cells with similar RNA profiles as a metacell. Specifically, we first selected non-M phase single cells from the embryo dataset, generating 3,217 cells in total. We normalized these cells using “SCTransform” by Seurat, performed PCA dimensionality reduction, and used the first 25 principal components at a resolution of 35 to cluster single cells into metacells. Metacells with fewer than 5 cells were discarded in subsequent analyses. We obtained a total of 167 metacells for the embryonic data, with each metacell consisting a median of 18 single cells. The metacell’s RNA profiles or compartments were generated by the mean of RNA expressions and scCompartments, for all single cells in this metacell.

For clustering of metacells, we used Seurat function “ScaleData” to normalize the metacell RNA profiles and performed PCA by “RunPCA” function with default parameters on top 2,000 variable features identified by “FindVariableFeatures” function. The cell typing of a metacell was carried out through a majority vote by all single cells within the metacell.

##### Pseudotime inference

For the HiRES data from developing mouse embryos, both the UMAP embedding of original scRNA-seq and metacell RNA-seq exhibited two differentiation trajectories, the neural trajectory (EN) and the mesenchymal trajectory (MLM), stemming from epiblast and primitive streak (EPI).

We performed pseudotime inference using Monocle3 Cao et al. (2019), based on the above UMAP embedding of metacells. We focused on the EN lineage and chose the EPI as starting points. Then we applied monocle3 to compute the pseudotime with default parameters.

### 2 Supplementary Results

#### 2.1 Simulation via downsampling high coverage scHi-C data

We then compared scDIAGRAM with 3D structure modeling from Hickit by downsampling the Dip-C dataset Tan et al. (2018), a high-coverage scHi-C dataset, using 14 GM12878 cells on chr1 at 500 kb resolution. The original data was downsampled from 50% to 5%. On the original data, the compartments generated by scDIAGRAM were consistent with the 3D structure modeling from Hickit (median intersection 0.85, Spearman correlation 0.7). scHiCluster produced a median intersection of 0.7 and a Spearman correlation of 0.45, with larger variance.

Upon downsampling, scDIAGRAM outperformed both scHiCluster and Higashi in terms of intersection and Spearman correlation (Supplementary Figure S18). At low sampling rates (below 10%), scDIAGRAM also outperformed Hickit (Supplementary Figure S18). Furthermore, across different sample rates, the intersection and correlation for scDIAGRAM barely decreased, demonstrating the robustness of our method to variations in sequencing depth.

In contrast, scHiCluster exhibited limited imputation benefits, with global Pearson and Spearman correlations always below 0.6. This suggests that 3D modeling and scHiCluster may generate quite different imputed matrices from the same scHi-C data (Supplementary Figure S18B), which limits their utility in certain applications. Higashi produced better imputation results than scHiCluster on this simulated dataset, as measured by global Spearman correlation; however, its performance still fell short of ideal (Supplementary Figure S18B).

### References


Abdennur, N., & Mirny, L. A. (2020). Cooler: scalable storage for Hi-C data and other genomically labeled arrays. *Bioinformatics*, *36*(1), 311–316. Retrieved from https://doi.org/10.1093/bioinformatics/btz540 DOI: 10.1093/bioinformatics/btz540

Bintu, B., Mateo, L. J., Su, J.-H., & et al. (2018). Super-resolution chromatin tracing reveals domains and cooperative interactions in single cells. *Science*, *362*, eaau1783.

Cao, J., Spielmann, M., Qiu, X., & et al. (2019). The single-cell transcriptional landscape of mammalian organogenesis. *Nature*, *566*, 496-502.

Gayoso, A., Shor, J., Carr, A. J., & et al. (2022). Jonathanshor/doubletdetection: doubletdetection v4.2. *Zenodo*. Retrieved from https://doi.org/10.5281/zenodo.6349517

Hafemeister, C., & Satija, R. (2019). Normalization and variance stabilization of single-cell Rna-seq data using regularized negative binomial regression. *Genome Biology*, *20*, 1-15.

Hao, Y., Hao, S., Andersen-Nissen, E., & et al. (2021). Integrated analysis of multimodal single-cell data. *Cell*, *184*, 3573-3587.e29.

Kent W.J, F. T., Sugnet CW., & et al. (2002). The Human Genome Browser at UCSC. *Genome Res.*, *12*(6), 996-1006.

Kim, H.-J., Yardımcı, G. G., Bonora, G., & et al. (2020). Capturing cell type-specific chromatin compartment patterns by applying topic modeling to single-cell Hi-C data. *PLoS Comput. Biol.*, *16*, e1008173.

Lee, D., Luo, C., Zhou, J., & et al. (2019). Simultaneous profiling of 3D genome structure and dna methylation in single human cells. *Nat Methods*, *16*, 999-1006.

Liu, Z., Chen, Y., Xia, Q., & et al. (2023). Linking genome structures to functions by simultaneous single-cell Hi-C and RNA-seq. *Science*, *380*, 1070-1076.

Nagano, T., Lubling, Y., Várnai, C., & et al. (2017). Cell-cycle dynamics of chromosomal organization at single-cell resolution. *Nature*, *547*, 61-67.

Newman, M. E. J. (2013). Spectral methods for community detection and graph partitioning. *Phys. Rev. E*, *88*, 042822.

Open2C, Abdennur, N., Abraham, S., & et al. (2024, 05). Cooltools: Enabling high-resolution Hi-C analysis in python. *PLOS Computational Biology*, *20*(5), 1-16. Retrieved from https://doi.org/10.1371/journal.pcbi.1012067

Ramani, V., Deng, X., Qiu, R., & et al. (2017). Massively multiplex single-cell Hi-C. *Nat. Methods*, *14*, 263-266.

Su, J.-H., Zheng, P., Kinrot, S. S., & et al. (2020). Genome-scale imaging of the 3D organization and transcriptional activity of chromatin. *Cell*, *182*, 1641-1659.

Tan, L., Xing, D., Chang, C.-H., Li, H., & Xie, X. S. (2018). Three-dimensional genome structures of single diploid human cells. *Science*, *361*, 924-928.

Van Dijk, D., & et al. (2018). Recovering gene interactions from single-cell data using data diffusion. *Cell*, *174*, 716-729.

Zhang, R., Zhou, T., & Ma, J. (2022). Multiscale and integrative single-cell Hi-C analysis with Higashi. *Nat. Biotechnol.*, *40*, 254-261.

Zhou, T., Zhang, R., Jia, D., & et al. (2024). GAGE-seq concurrently profiles multiscale 3D genome organization and gene expression in single cells. *Nat. Genet.*, *56*, 1701-1711.

#### 2.2 Supplementary Figures

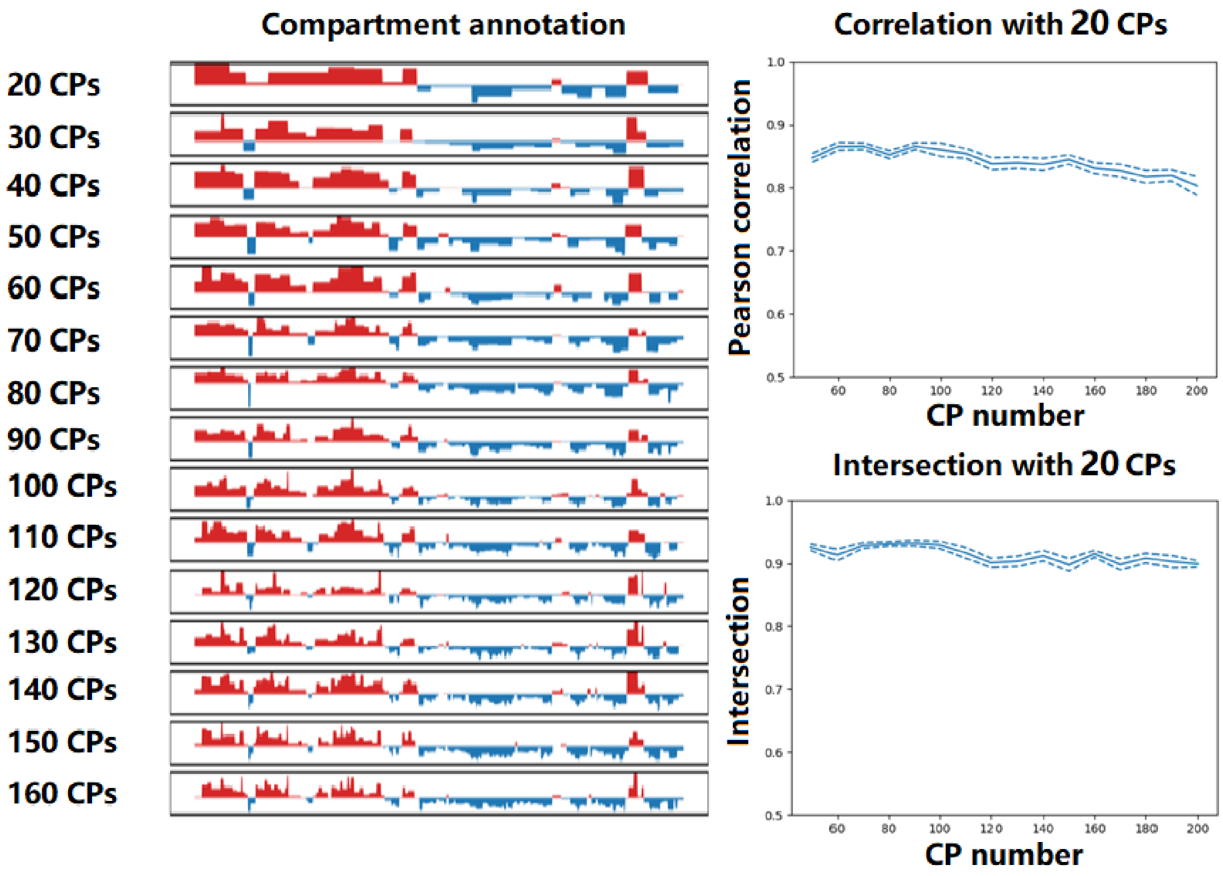


**Supplementary Fig. S1: Robustness of scDIAGRAM to the choice of CP number on a real single cell (GAGE-seq, chr1, 100kb).** scDIAGRAM exhibited robustness to CP number selection. Testing on a single cell (GAGE-seq, chr1 100kb pre-centromere, not many CPs in bulk PCA), results stabilized at 20 CPs. Further increases minimally affected output, as seen in visual concordance and high correlation/intersection with the 20-CP results.

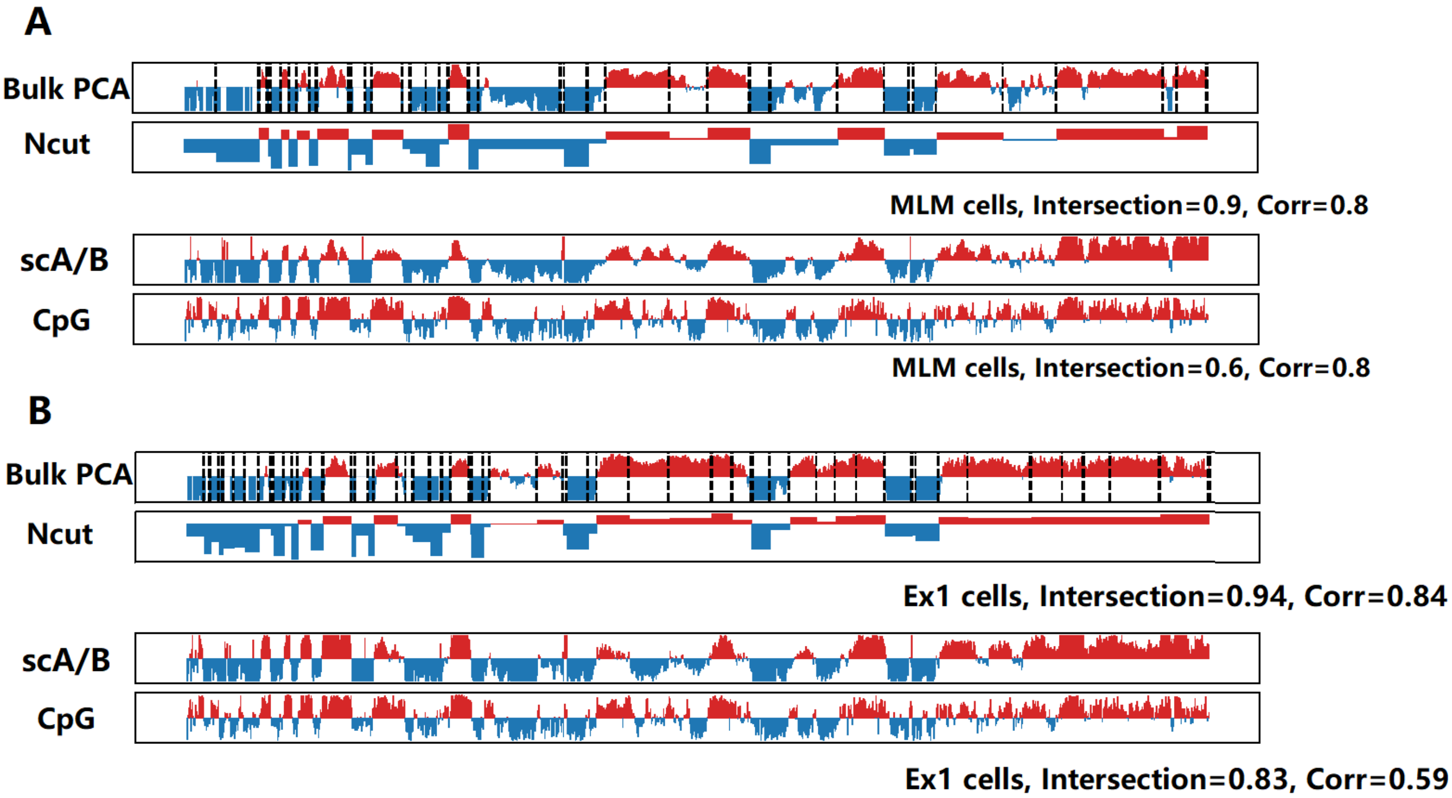


**Supplementary Fig. S2. Compartment annotation of pseudo-bulk Hi-C (100kb, chr7) in mix late mesenchyme (MLM) and Ex1 cell types.** (A) From MLM pseudo-bulk Hi-C we computed bulk PCA (40 CPs, dashed lines), Ncut, scA/B, and CpG density (from the reference genome), with intersections/Pearson correlations between both Ncut and scA/B with bulk PCA. (B) Same analyses as in (A), using the Ex1 pseudo-bulk Hi-C matrix.

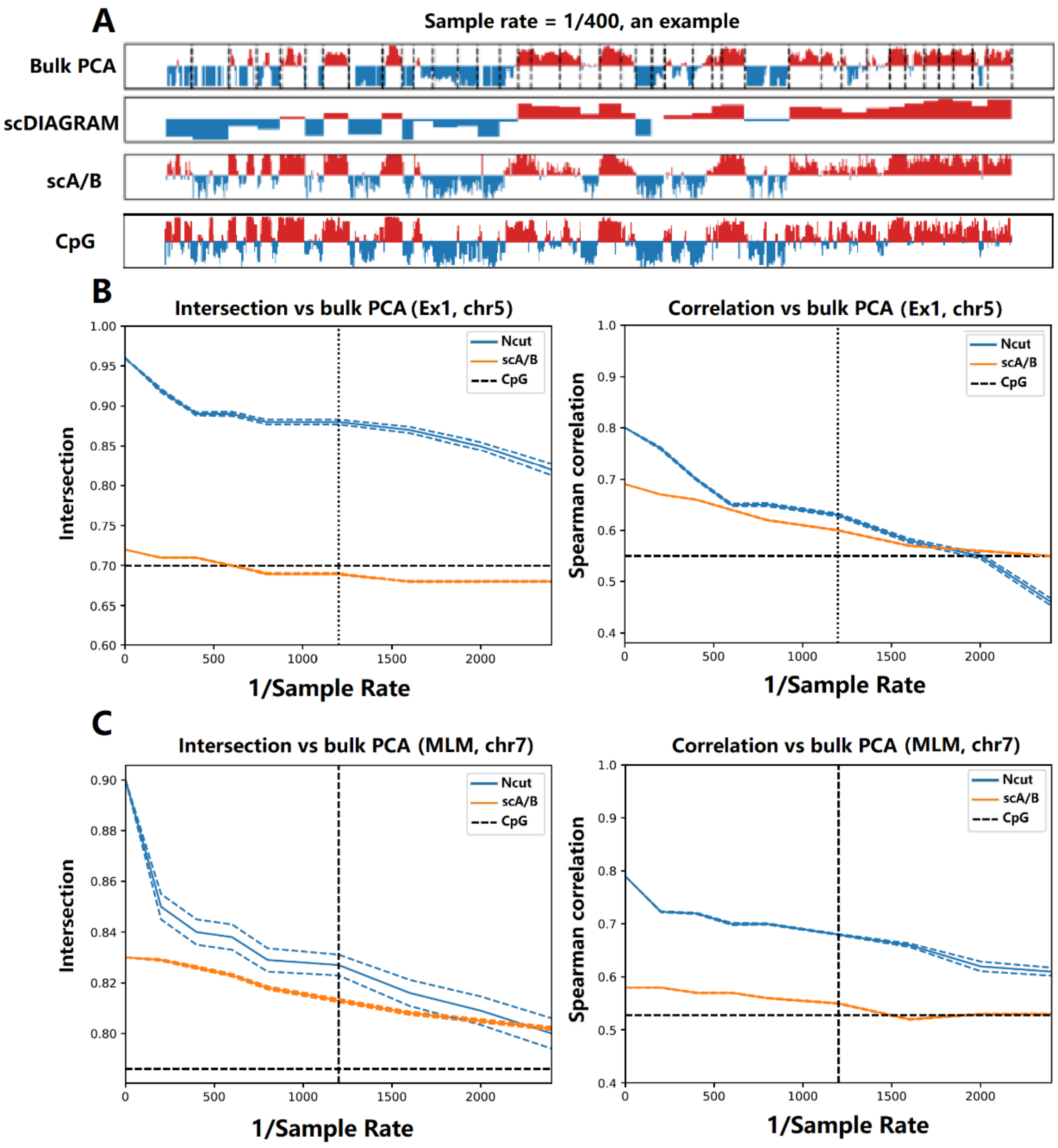


**Supplementary Fig. S3. Performance evaluation of scDIAGRAM on downsampled pseudo-bulk Hi-C data.** (A) For Ex1 pseudo-bulk Hi-C at 100kb resolution (1/400 sampling rate), we compared compartments from scDIAGRAM and scA/B against the ground-truth bulk PCA (40 CPs), along with CpG density profiles. (B-C) Intersection and Spearmans correlation for downsampled data from the (B) Ex1 (chr5) and (C) MLM (chr7) cell type, with vertical dashed lines indicating real scHi-C sampling rates and horizontal lines showing CpG vs bulk PCA intersection/correlation levels. At the real scHi-C sampling rate 1/1200, scDIAGRAM was always better than scA/B. At the rate 1/2400, when the downsampled Hi-C matrices was too sparse, scDIAGRAM might be worse than scA/B as scA/B would approximate the CpG level.

| Datasets | Contact numbers per cell | | | |
|---|---|---|---|---|
| | Mean | Min | Median | Max |
| Nagano et al.(2017) | 4.4 k | 0.2 k | 4.4 k | 19.4 k |
| WTC-11 (Zhang et al. (2022)) | 14.5 k | 2.2 k | 14.7 k | 25.3 k |
| Ramani et al. (2017) | 0.9 k | 0.0 k | 0.7 k | 9.5 k |
| 4DN sci-Hi-C (Kim et al. (2020)) | 0.6 k | 0.0 k | 0.4 k | 77.3 k |
| sn-m3c-seq (Lee et al. (2019)) | 14.0 k | 0.4 k | 15.5 k | 37.9 k |
| HiRES (Liu et al. (2023)) | 24.4 k | 4 k | 23.6 k | 42.8 k |
| GAGE-seq (Zhou et al. (2024)) | 26.6 k | 0.1 k | 23.4 k | 202.8 k |
| Dip-C GM12878 (Tan et al. (2018)) | 43.9 k | 32.2 k | 43.4 k | 67.5 k |

**Supplementary Figure S4. Statistics of the contact numbers per cell at 1Mb resolution of chr2.** Statistics of the contact numbers per cell at 1Mb resolution of chr2 for different datasets. The Nagano et al. data were mapped to the mm9 assembly. The Ramani et al. and sn-m3c-seq data were mapped to the hg19 assembly. HiRES and GAGE-seq were mapped to the mm10 assembly. Others were mapped to the hg38 assembly.

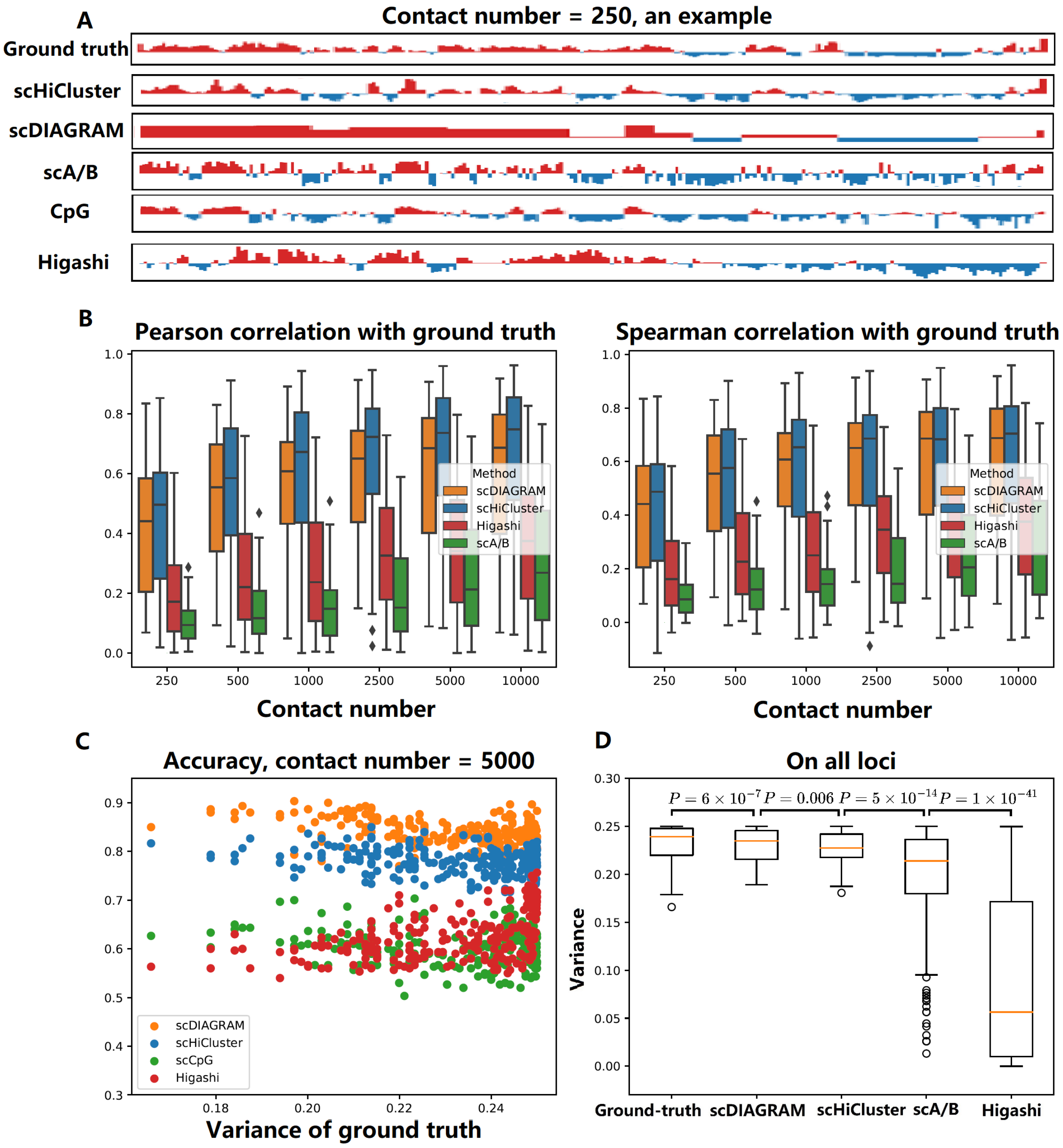


**Supplementary Fig. S5. Performance on downsampled imaging data (Su et al. IMR-90, chr2).** (A) At 1Mb resolution (downsampled with 250 contacts), we showed the ground-truth bulk PCA compared with scHiCluster, scDIAGRAM, scA/B compartments and the CpG density. (B) Correlation analysis revealed scDIAGRAM performed comparably to scHiCluster, while scA/B showed the lowest agreement with bulk PCA. Higashi performed slightly better than scA/B, but inferior to scDIAGRAM and scHiCluster. (C) When downsampled at 5000 contacts, scDIAGRAM maintained the highest accuracy versus other methods. (D) scDIAGRAM exhibited greater heterogeneity than scHiCluster, Higashi and scA/B across all loci.

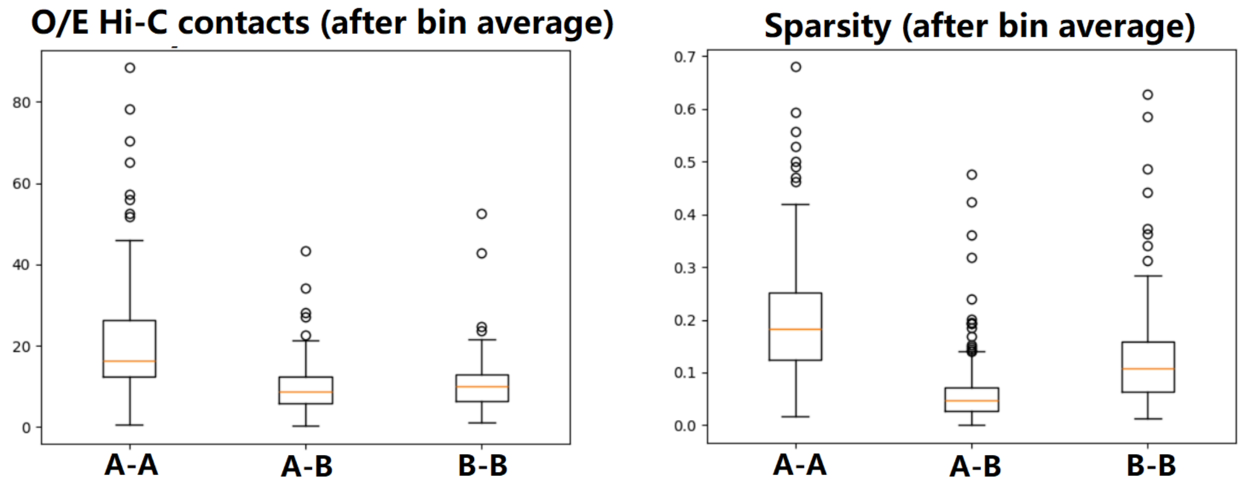


**Supplementary Fig. S6. Compartment analysis on human GM12878 cells.** Median observed-over-expected (O/E) contact frequencies (A-A, A-B, B-B) from scDIAGRAM after bin averaging, with corresponding contact sparsities. The median was computed for nonzero values.

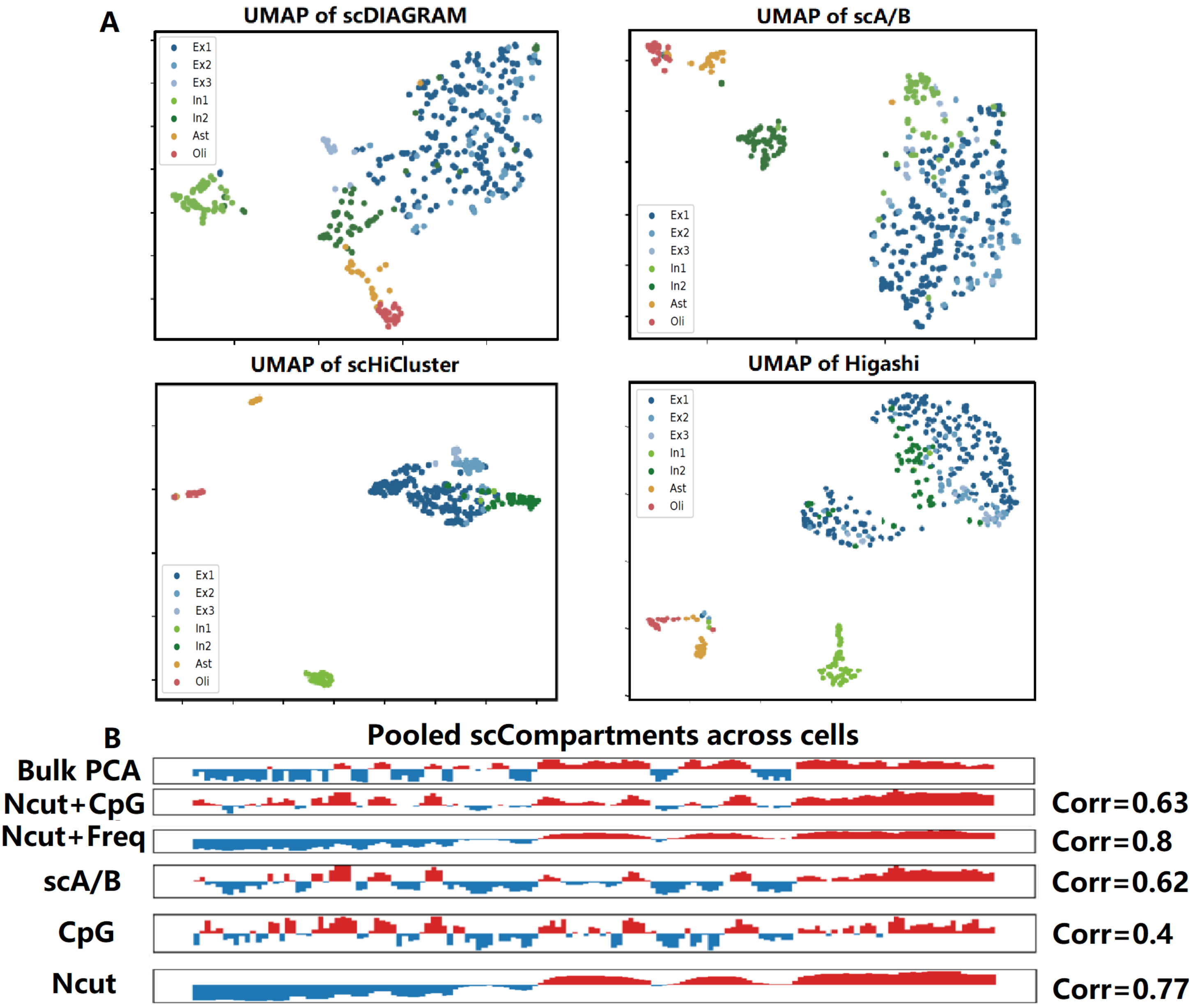


**Supplementary Fig. S7. Embedding and Pooled compartments of the mouse brain neurons.** (A) UMAP embeddings of single cells using scDIAGRAM, scA/B, Higashi and scHiCluster, with scRNA-seq annotations (Ex1-3: excitatory neurons; In1-2: inhibitory neurons; Ast: astrocyte; Oli: oligodendrocyte). scDIAGRAM, scHiCluster and Higashi showed similar clustering patterns (In2 was more close to Ex1-3), while scA/B exhibited distinct patterns. (B) Pooled Ex1 scCompartments (chr7, 1Mb) compared to bulk PCA, with Pearson correlations calculated. Ncut, Ncut+Freq and Ncut+CpG compartments were studied alongside scA/B and CpG. Pooled scDIAGRAM (Ncut) correlated better with bulk PCA than scA/B. When combining Ncut with CpG, unlike in other analysis, it would reduce the correlation.

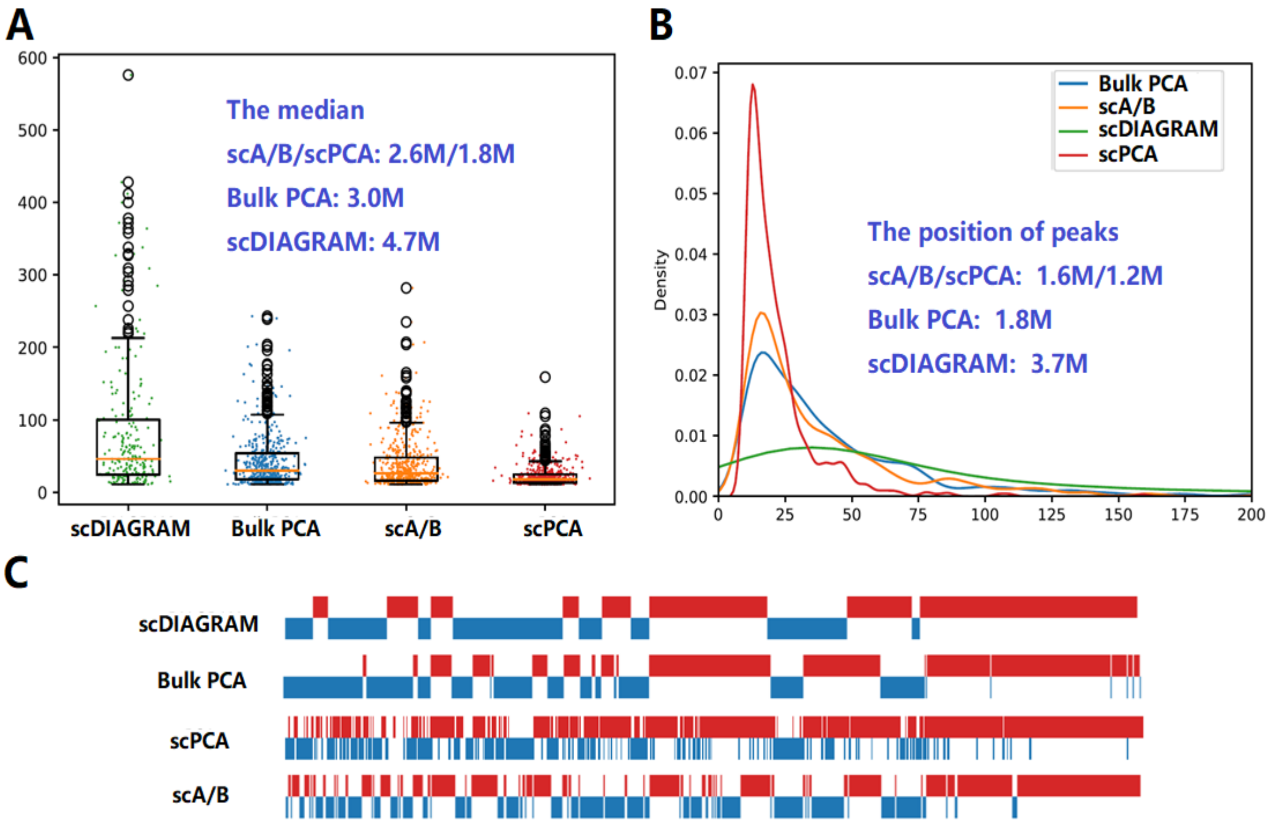


**Supplementary Fig. S8. Compartmental length of the mouse brain neurons**. Results were generated from one cell on the whole genome at 1 Mb resolution. (A) Boxplot comparing compartment lengths across methods, with scDIAGRAM showing closest median length to bulk PCA. (B) Unimodal kernel density distributions revealed scDIAGRAMs peak position most closely matched bulk PCA. (C) Example binary compartments demonstrating compartmental lengths for each method.

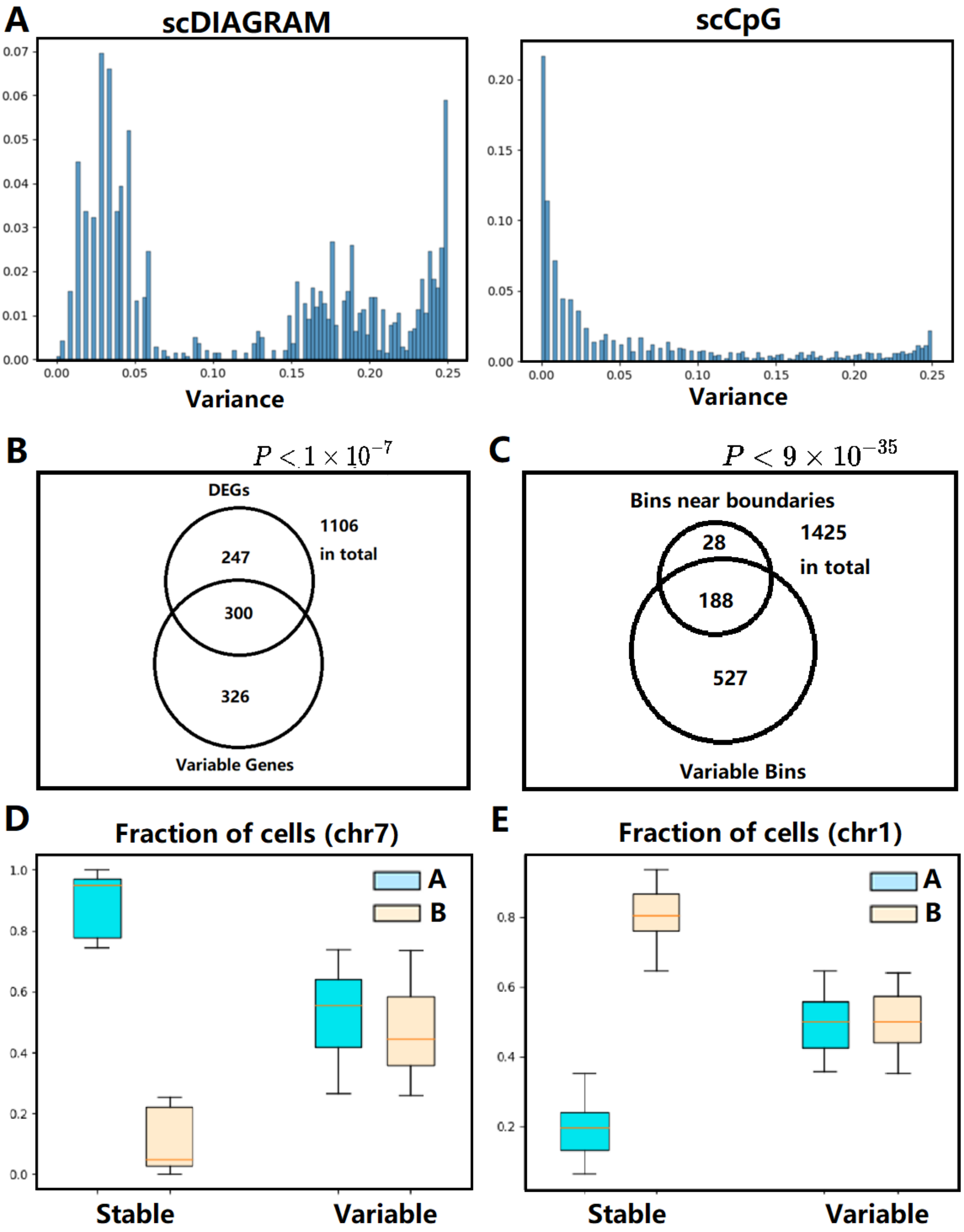


**Supplementary Fig. S9. Stable/variable bins of the mouse brain neuron scHi-C data.** (A) Bimodal variance distributions (scDIAGRAM and scA/B) revealed stable and variable genomic loci. (B) Compartment variability (for different cell types, across the dataset) correlated with cell-type-specific marker genes. (C) Within Ex1 cells, the compartmental variability associated with compartmental boundaries. (D-E) Fraction of cells for stable/variable bins in A/B compartments, on different chromosomes. Variable regions distributed uniformly across A/B compartments, while stable regions predominantly occupy a single compartment. Here we used the top and lower 25-th percentile as the threshold to determine stable/variable regions.

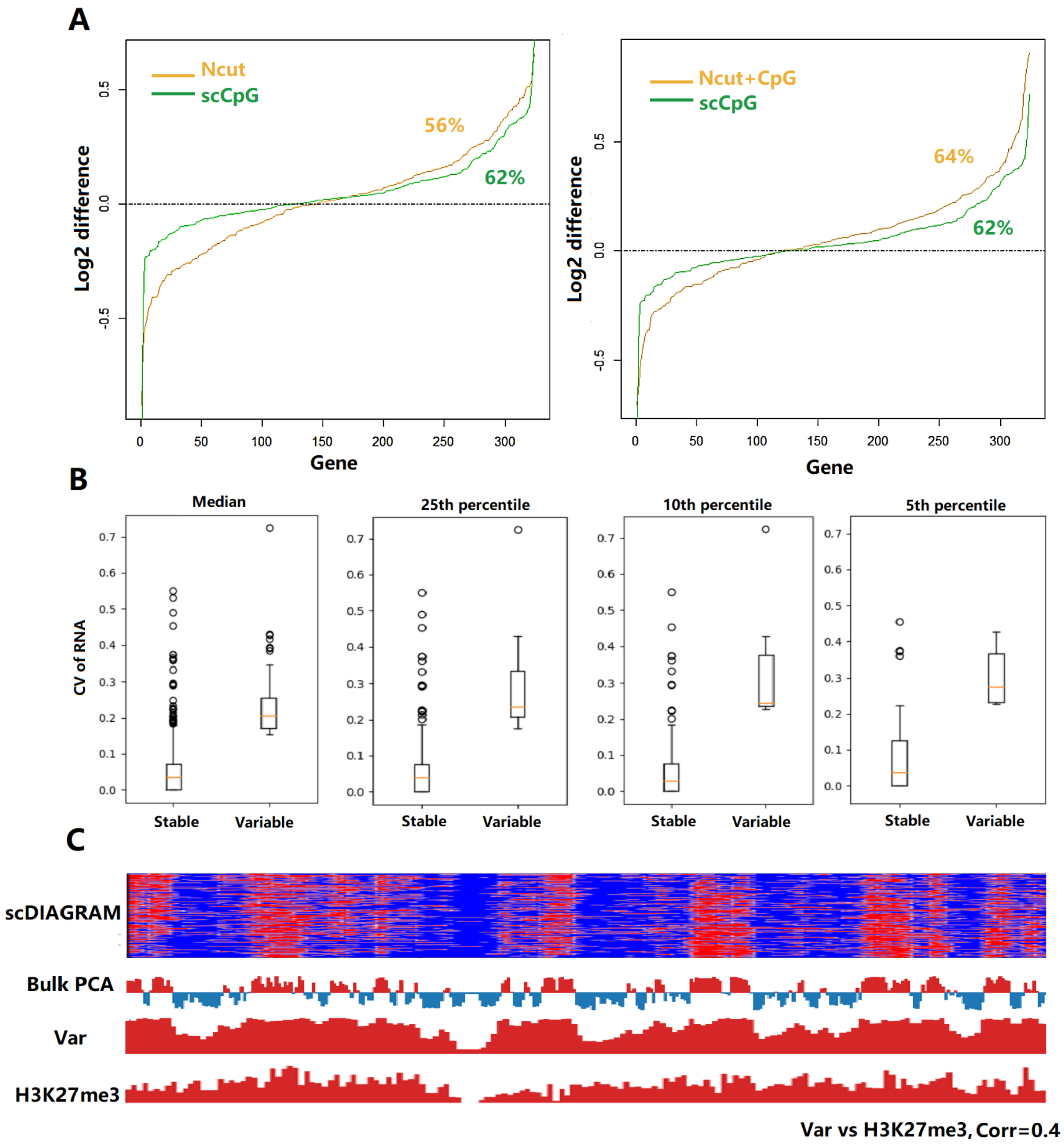


**Supplementary Fig. S10. Linking single-cell compartments with RNA expressions in mouse brain neurons.** (A) In Ex1 cells, transcribing genes (UMI >10) showed more active compartments than silent genes (UMI <1). We took average of scCompartments across these states after normalized into [0,1], then we computed the log2 difference. We also computed the ratio of genes that was compartmental more activated in transcribing states. The ratio was comparable among these methods. Ncut typically generated larger compartmental difference compared with scA/B, indicating scDIAGRAM was more heterogenous. (B) The robustness of our results when different thresholds were used. We computed the transcriptional variability (CV) for variable/stable regions, when different thresholds were used to determine the variable/stable groups. (C) In GM12878 cells (chr7, 1Mb), we showed the heatmap from scDIAGRAM and bulk PCA. Then we found scDIAGRAM's variance was correlated with H3K27me3, a mark enriched in B1 subcompartments and hence indicated the region's compartmental instability.

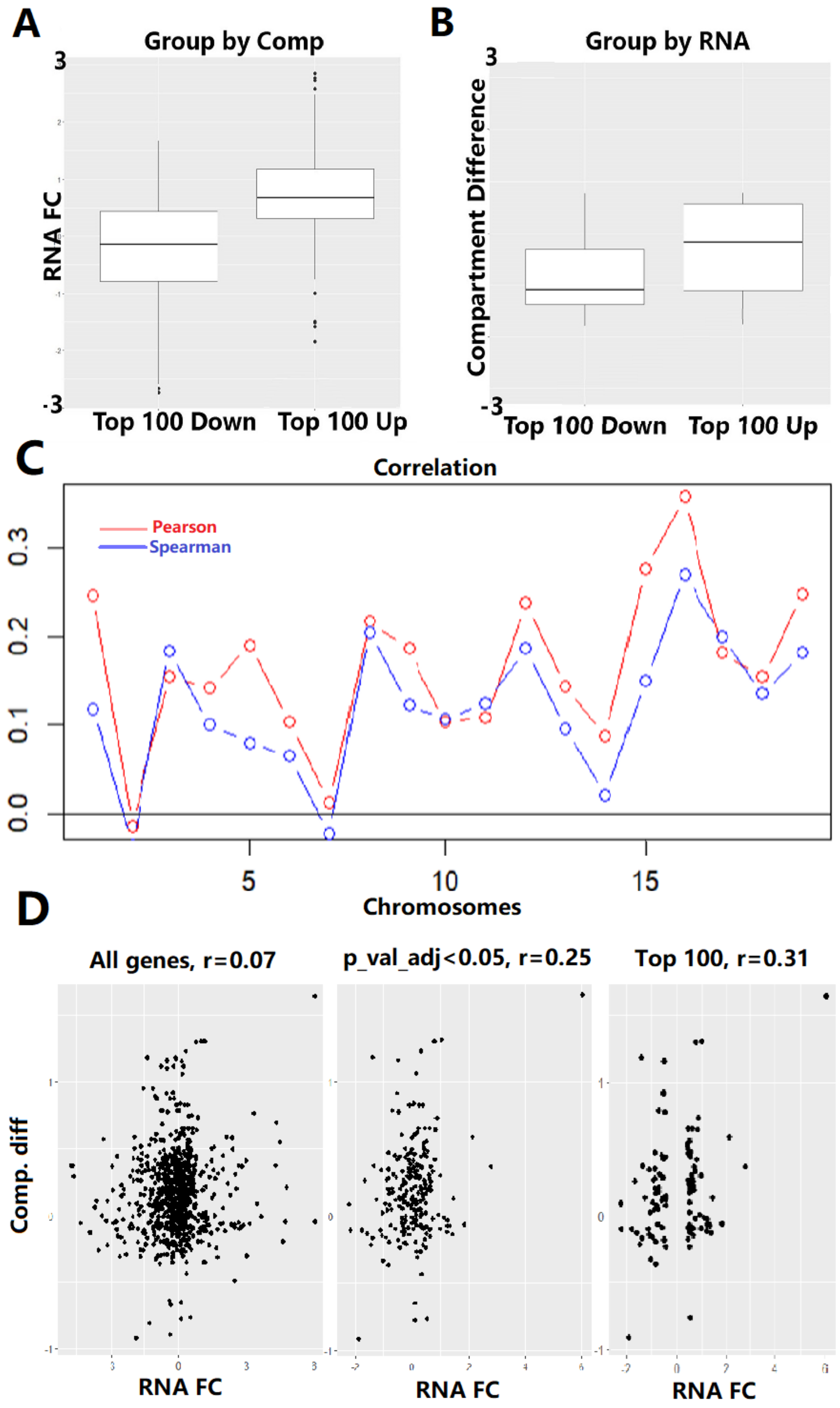


**Supplementary Fig. S11. Compartment-RNA relationships between two cell types, using mouse brain neurons.** (A-B) Comparing the transcription and compartmental transition between Ex1 and Ast cells: (A) Top 100 compartment-activated genes (by scDIAGRAM) showed higher RNA upregulation than top 100 inactivated genes, while (B) top 100 up-regulated genes exhibited compartmental activation than top 100 down-regulated genes, suggesting compartment changes may influence transcription. In (A) we observed larger difference between two groups, indicating compartmental changes as a potential cause for RNA transcription. (C) Chromosome-wide analysis revealed positive RNA-compartment correlations (comparing RNA fold change with compartmental differences) between excitatory/inhibitory neurons, except for chr2/chr7. (D) In chr1, stronger correlations emerged with increasingly stringent gene selection: all genes expressed in at least 1% cells (760 genes) → significant genes (230 genes, adjusted $p<0.05$) → top 100 genes ordered by fold-change, highlighting compartment-RNA coupling.

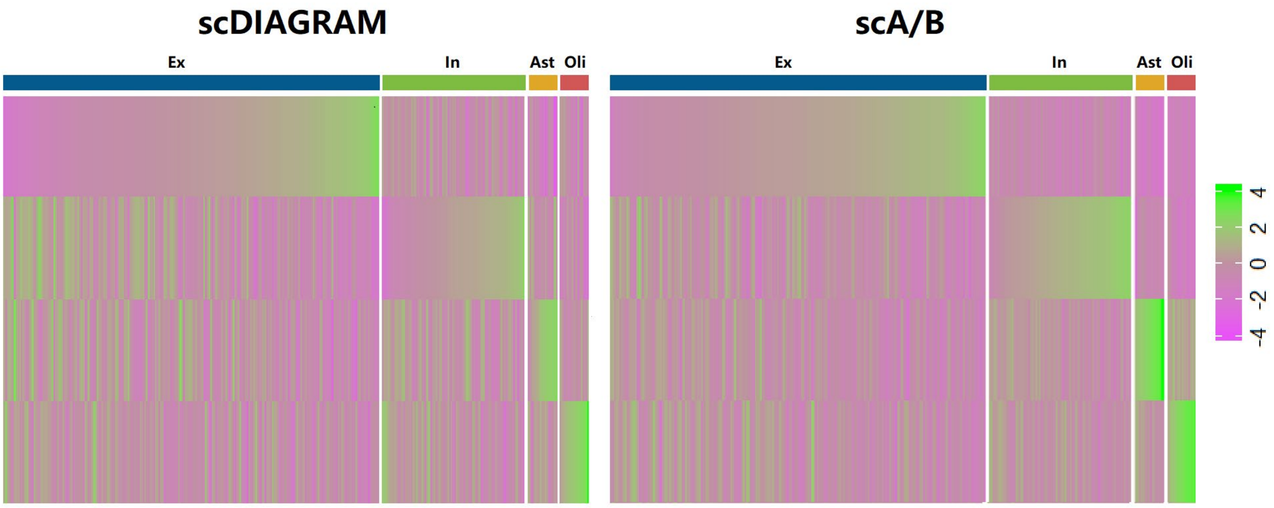


**Supplementary Fig. S12. Compartment values on cell-type-specific marker genes (mouse brain neurons).** Mean compartmental values for top 500 marker genes across 4 cell types. Each column referred to a single cell (ordered by their cell types and the compartmental enrichment at the corresponding markers). Each row referred to a marker gene set. We used real-valued compartments from scDIAGRAM and each row was z-score normalized after averaging across markers.

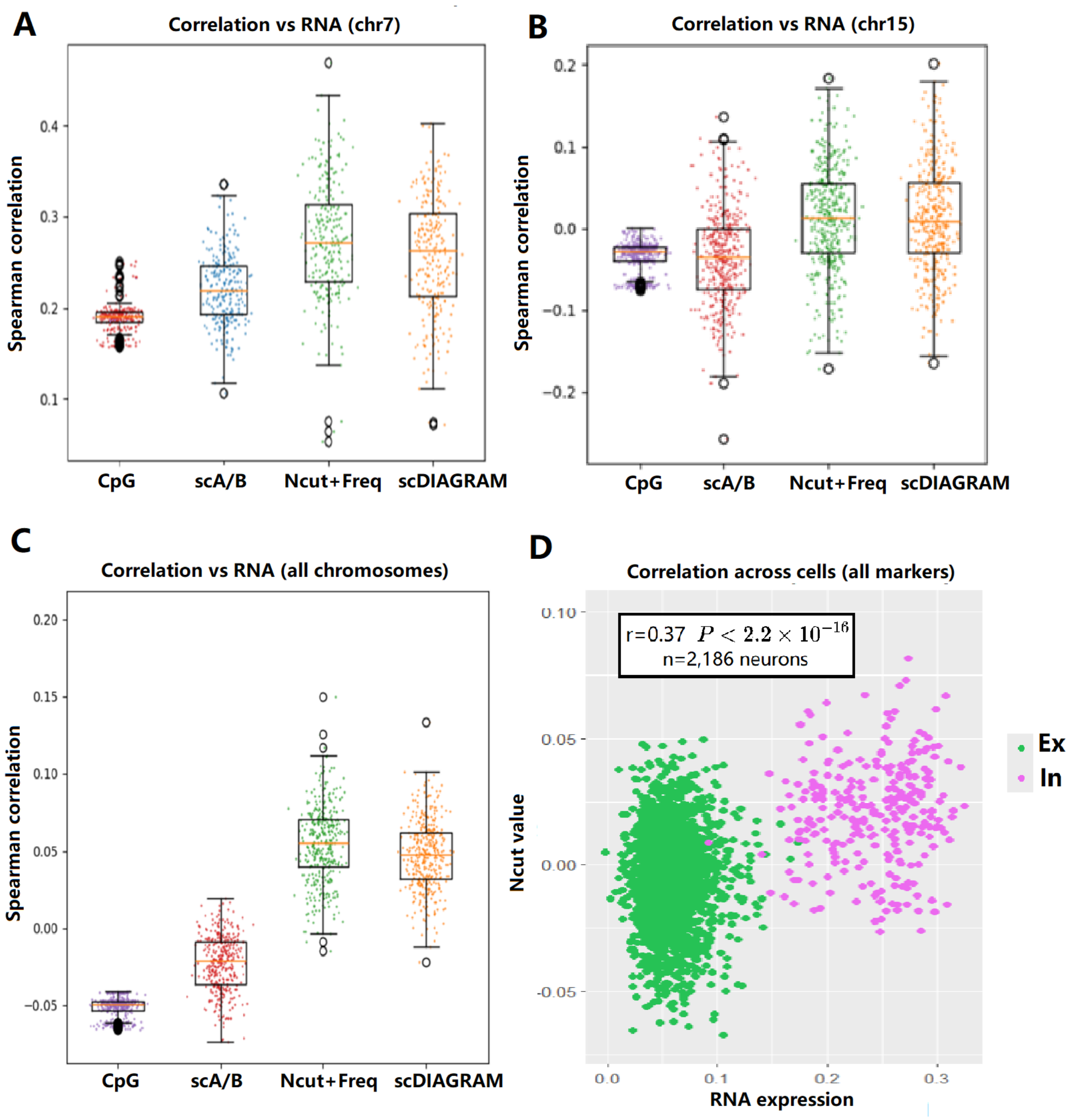


**Supplementary Fig. S13. Compartment-RNA correlations in mouse brain neurons.** (A-B) Spearman correlations between RNA expressions and real-valued compartments from scDIAGRAM, on chr7 and chr15. (C) Spearman correlations for all chromosomes. (A-C) used the HiRES dataset. (D) Significant positive correlation (r=0.37) between scDIAGRAM compartments and marker gene expression (n=2,186 Ex vs. In neurons, using the GAGE-seq dataset).

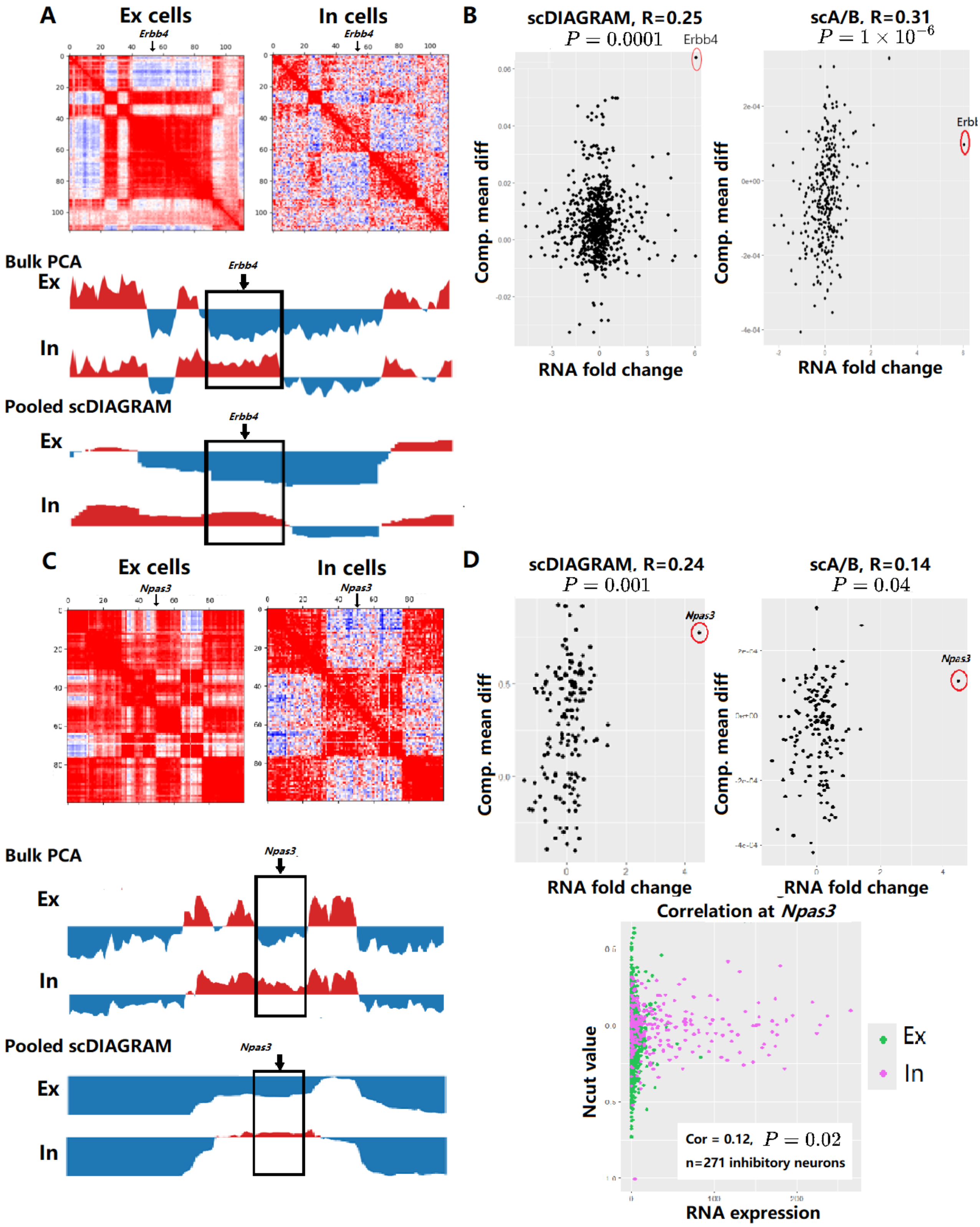


**Supplementary Fig. S14. Compartment-RNA relationships at *Erbb4* and *Npas3* loci.** (A,C) Pseudo-bulk Hi-C matrices, bulk PCA, and pooled scDIAGRAM compartments at (A) *Erbb4* and (C) *Npas3* loci. (B,D) Pearson's correlations between compartment differences (for scDIAGRAM and scA/B) and RNA log fold changes (In vs Ex) for (B) chr1 DEGs (n=230 genes, one-sided tests for nonzero correlations) and (D) chr12 DEGs (n=158 genes), with

*Erbb4* and *Npas3* showing the most significant increase in both RNA expressions and compartments (by scDIAGRAM). scA/B only exhibited moderate difference on these two loci, indicating scDIAGRAM was more consistent with RNA observations. In (D) we also showed the correlation between RNA expressions and real-valued scCompartments (from scDIAGRAM) at this locus across single cells.

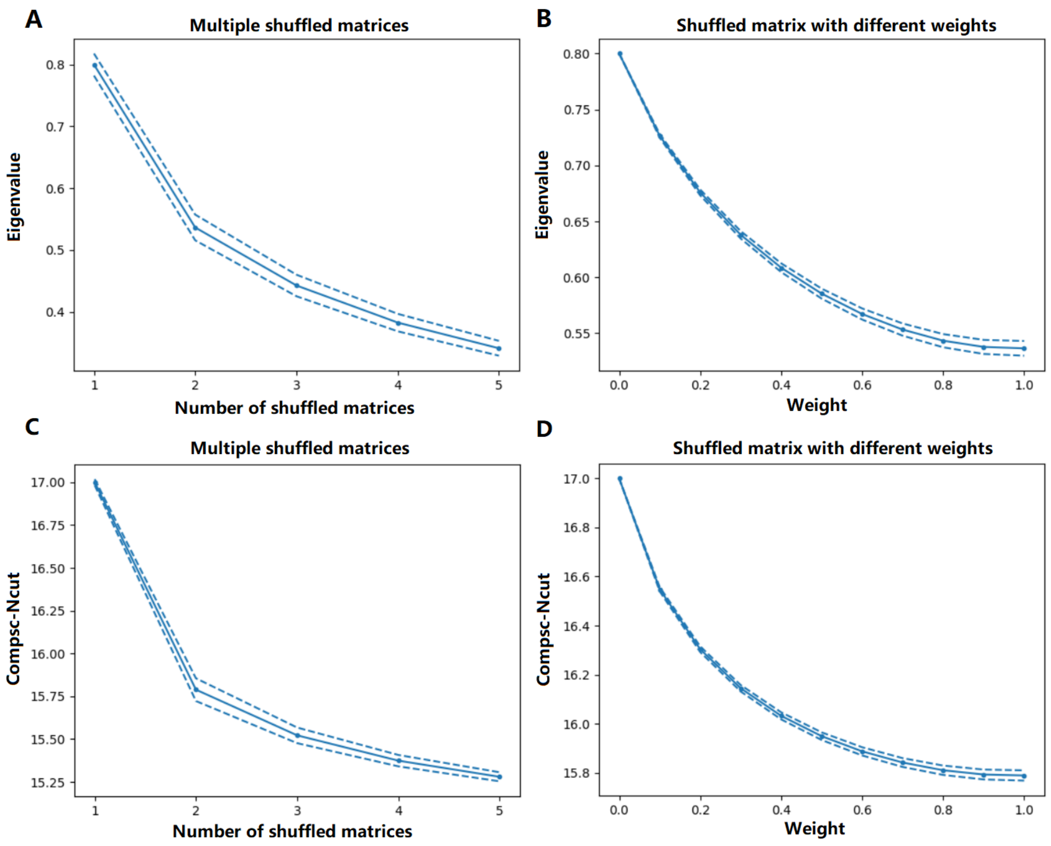


**Supplementary Fig. S15. The eigenvalue and Compsc-Ncut in the permutation experiments.** (A-B) The second eigenvalue changes when: (A) pooling original data with K=1-5 shuffled matrices, or (B) pooling original data with a single weighted shuffled matrix (0.1-0.9 weights). (C-D) Corresponding Compsc-Ncut changes under the same conditions.

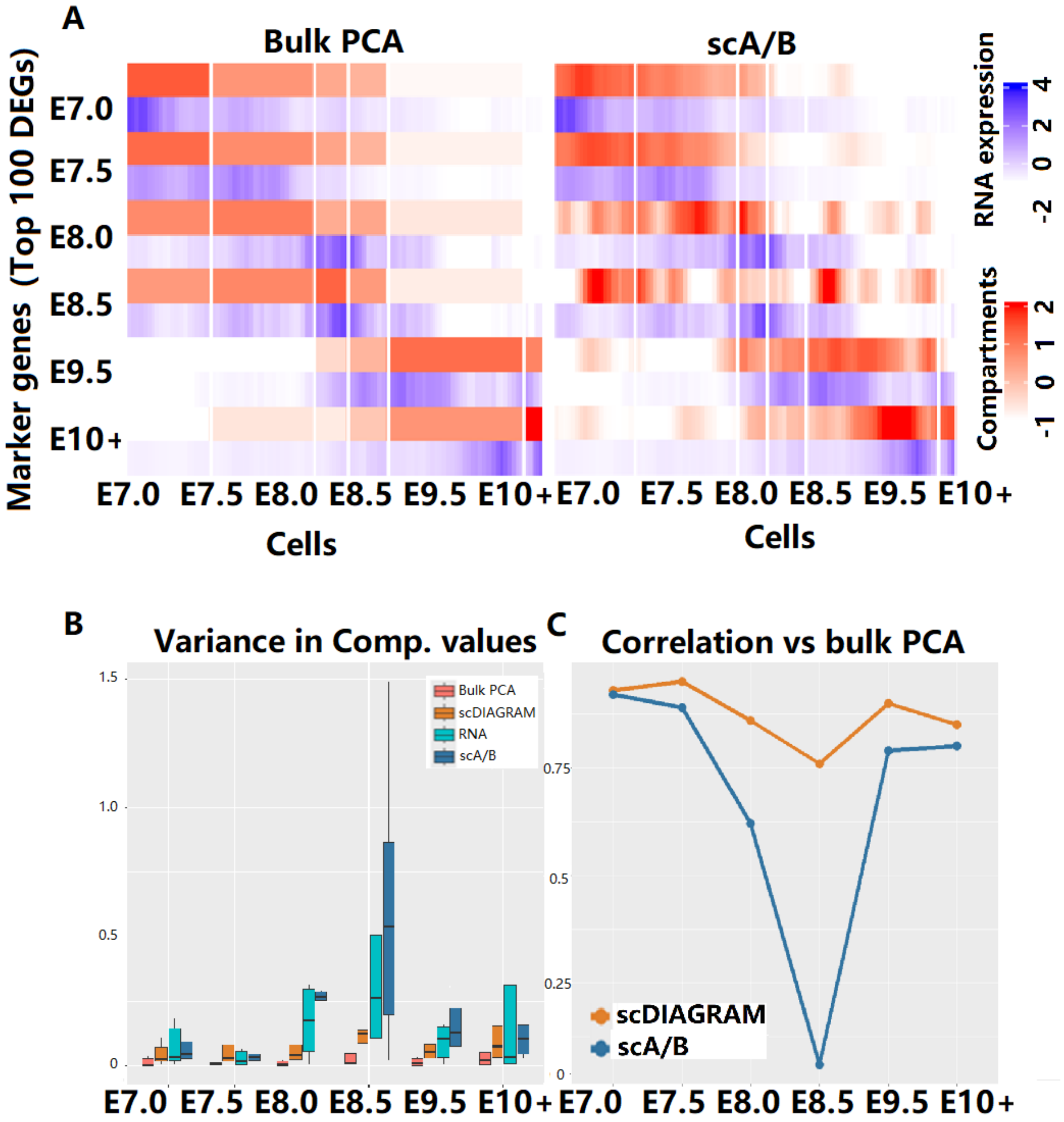


**Supplementary Fig. S16. Compartment dynamics during embryo development (E7.0-E10+).** (A) Stage-specific marker gene enrichment (pseudo-time ordered) of compartments from bulk PCA and scA/B. Bulk PCA was calculated from pseudo-bulk matrices at each stage. The top 100 marker genes (ordered by fold change) at each stage were used. (B) Compartmental variances at each stage for different methods. (C) scDIAGRAM correlated more closely with bulk PCA than scA/B (row-wise correlations for heatmaps in (A) and Fig. 4C in the maintext).

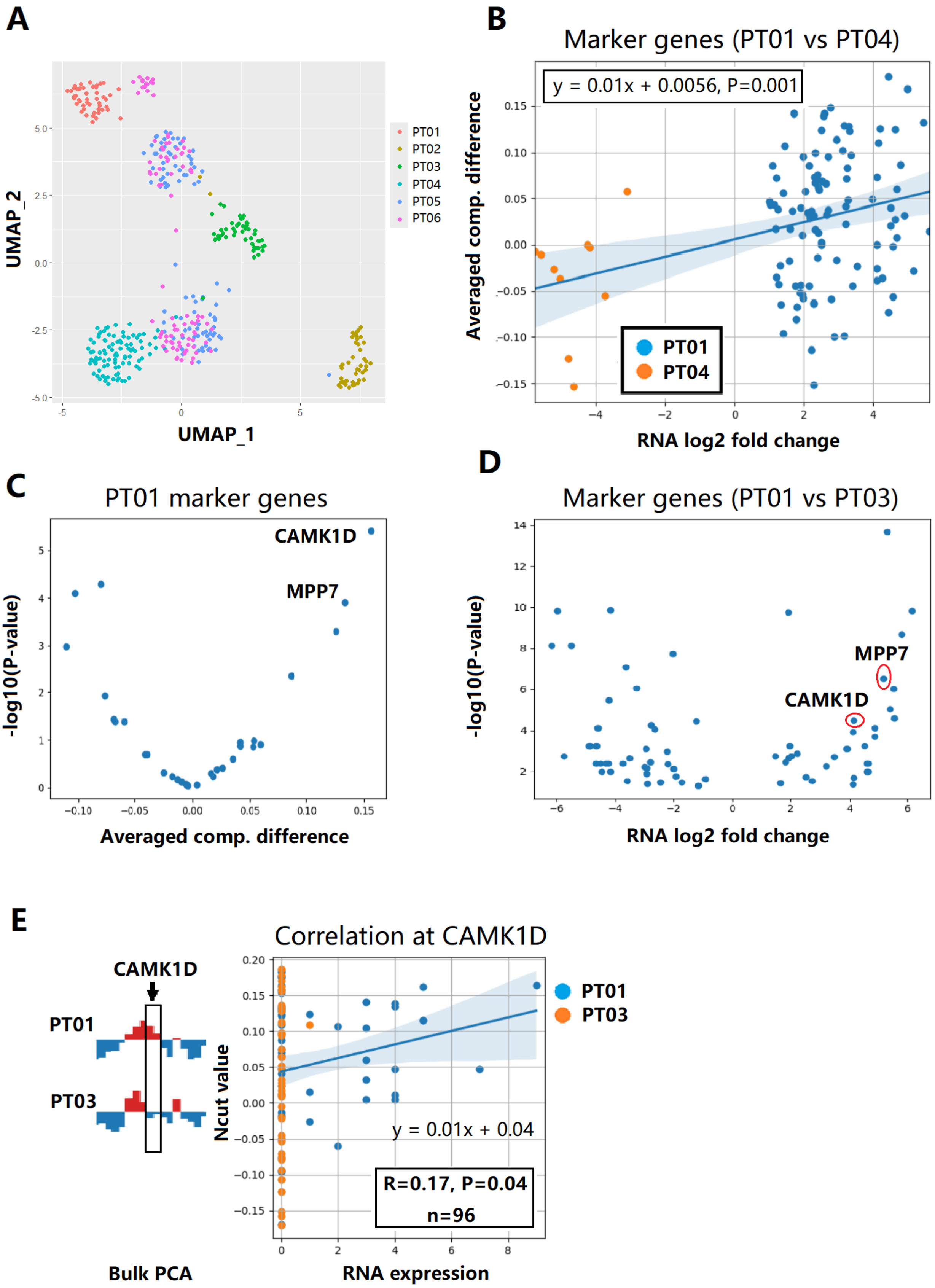


**Supplementary Fig. S17. Compartmental differences in AML.** (A) UMAP embedding of scRNA-seq data showed patient-specific clustering, motivating our focus on compartmental differences between patients. (B) Correlation between changes in compartmentalization and gene expression for marker genes comparing patients PT01 and PT04. (C) Volcano plot showing differential scDIAGRAM compartment values between PT01 and PT03. P-values were calculated using a two-sample t-test. Only marker genes from PT01 are shown. (D) Volcano plot of differential gene expression between PT01 and PT03, with P-values computed using the MAST test in Seurat. (E) Visualization of bulk PCA signals at the CAMK1D locus in PT01 and PT03, along with RNA expression and scDIAGRAM Ncut values for each single cell at the same locus.

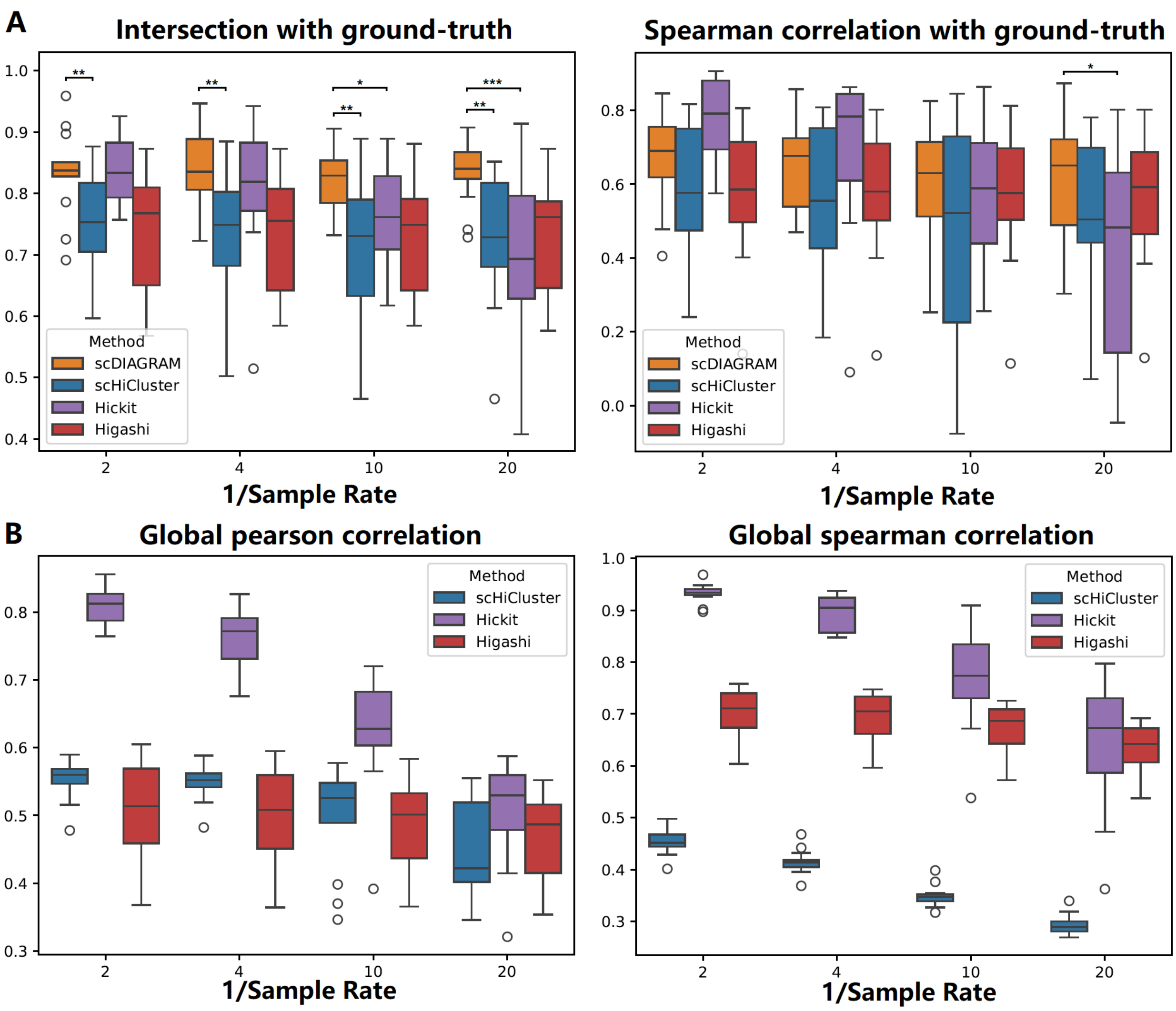


**Supplementary Fig. S18. Performance on downsampled high-coverage scHi-C (the DipC data).** (A) Using Hickit (rate=1) as ground truth (n=14 GM12878 cells in total, chr1, at 500kb resolution), scDIAGRAM outperformed scHiCluster and Higashi at all sampling rates. It performed better than Hickit at lower rates where Hickit performance declined. (B) Global Pearson/Spearman correlations (the correlation between two flattened Hi-C matrices) with the ground-truth showed imputation methods (scHiCluster, Higashi, Hickit) produced different but increasingly similar imputation matrices when the sample rate decreased. Higashi demonstrating the greatest robustness to downsampling.
